# Comparative single-cell analysis reveals IFN-γ as a driver of respiratory sequelae post COVID-19

**DOI:** 10.1101/2023.10.03.560739

**Authors:** Chaofan Li, Wei Qian, Xiaoqin Wei, Harish Narasimhan, Yue Wu, Mohd Arish, In Su Cheon, Kamya Sharifi, Ryan Kern, Robert Vassallo, Jie Sun

## Abstract

Post-acute sequelae of SARS-CoV-2 infection (PASC) represents an urgent public health challenge, with its impact resonating in over 60 million individuals globally. While a growing body of evidence suggests that dysregulated immune reactions may be linked with PASC symptoms, most investigations have primarily centered around blood studies, with few focusing on samples derived from post-COVID affected tissues. Further, clinical studies alone often provide correlative insights rather than causal relationships. Thus, it is essential to compare clinical samples with relevant animal models and conduct functional experiments to truly understand the etiology of PASC. In this study, we have made comprehensive comparisons between bronchoalveolar lavage fluid (BAL) single-cell RNA sequencing (scRNAseq) data derived from clinical PASC samples and relevant PASC mouse models. This revealed a strong pro-fibrotic monocyte-derived macrophage response in respiratory PASC (R-PASC) in both humans and mice, and abnormal interactions between pulmonary macrophages and respiratory resident T cells. IFN-γ emerged as a key node mediating the immune anomalies in R-PASC. Strikingly, neutralizing IFN-γ post the resolution of acute infection reduced lung inflammation, tissue fibrosis, and improved pulmonary gas-exchange function in two mouse models of R-PASC. Our study underscores the importance of performing comparative analysis to understand the root cause of PASC for developing effective therapies.

## Introduction

Three years after the onset of the SARS-CoV-2 pandemic, the development of anti-viral therapies and effective vaccines have significantly improved the management of acute COVID-19. However, an urgent public health challenge remaining is the presence of an increasingly large number of individuals (estimated to be more than 60 million globally) with post-acute sequelae of SARS-CoV-2 infection (PASC)^1^, ^2^. PASC are characterized by persistent, recurring, or novel symptoms manifesting after the resolution of acute infection^3^. Longitudinal assessments have indicated that PASC contributed 80 – 642 disability-adjusted life years (DALYs) per 1,000 individuals recovering from COVID-19, with over 20% experiencing symptoms beyond two years^2^. As the lung is the primarily affected organ during acute infection, respiratory PASC (R-PASC), which includes disabling symptoms like dyspnea, cough, and interstitial lung disease, is recognized as one of the predominant sequelae^3, 4, 5^. Furthermore, impaired gas-exchange functions may lead to systemic symptoms, including exertional dyspnea, chronic fatigue, and other limitations due to chronic hypoxia. Of note, respiratory symptoms, impairment in lung gas exchange, and radiographic abnormalities can persist for more than 2 years (possibly longer) in some patients, especially following severe acute COVID-19 infection^6, 7, 8^.

Emerging evidence has associated prolonged or aberrant peripheral immune responses with systemic or multi-organ symptoms observed in PASC, mainly through analyses of PBMCs and plasma^9, 10, 11, 12^. Currently, the immune status within affected organs during PASCs remains largely uncharacterized, mainly due to the limited direct sampling of affected tissues after acute COVID-19. Furthermore, it is extremely challenging to determine the causative mechanisms of PASC solely based on observational clinical research, as most clinical studies can only provide correlative phenotypic associations. Notably, a few animal models on COVID-19 sequelae have been developed^13, 14^. However, whether these animal models can faithfully model the pathophysiology and molecular etiology of human post COVID-19 sequelae remains to be determined since there is a lack of direct comparative analysis of these models with human investigations. Thus, there is an urgent need to perform a parallel comprehensive analysis of clinical samples and samples from relevant animal models side-by-side to acquire deeper mechanistic insights regarding PASC etiology, particularly in the respiratory tract.

Additionally, the development of future therapeutic interventions necessitates functional studies in clinically relevant animal models for validating the causative drivers of PASC. To address these challenges, here we undertook single-cell RNA (scRNAseq) sequencing of bronchoalveolar lavage (BAL) cells and peripheral blood mononuclear cells (PBMCs) from a cohort of COVID-19 convalescents with or without respiratory PASC (R-PASC). In parallel, we have performed kinetical scRNAseq analysis of BAL samples obtained from SARS-CoV-2 mouse models. The comparative scRNAseq analyses of human and animal samples revealed that R-PASC are associated with aberrant responses and interactions of macrophages and resident T cells in the respiratory tract. We further discovered IFN-γ as a central node mediating the aberrant interactions between respiratory macrophages and T cells. Strikingly, neutralization of IFN-γ post the resolution of acute viral infection dampened chronic pulmonary inflammation and tissue fibrosis, as well as restored pulmonary gas-exchange function in mouse R-PASC models. Our findings thus open the door for therapeutic interventions against R-PASC by targeting aberrant respiratory immunity in COVID-19 convalescents, especially IFN-γ.

## Results

### Single-Cell RNA-Seq reveals altered pulmonary immune landscape in R-PASC

The comprehensive respiratory immune and molecular landscape in COVID-19 convalescents, particularly in R-PASC, have not been characterized. To this end, we utilized a cohort of COVID-19 convalescents and controls that were not previously infected with SARS-CoV-2 (non-COVID-19). All convalescents were discharged and evaluated 60-90 days after the onset of SARS-CoV-2 infection and were SARS-CoV-2 N1 gene PCR negative prior to recruitment, as detailed previously^15^ (Fig. 1a). Pulmonary function was assessed using spirometry and lung diffusion capacity tests, including FVC (forced vital capacity), indicative of total lung capacity; FEV1 (forced expiratory volume in one second), indicative of lung volume exhaled in one second as a measure of obstructive airflow; and DLCO (Diffusing capacity for carbon monoxide), indicative of lung gas exchange efficacy.

**Figure 1.**
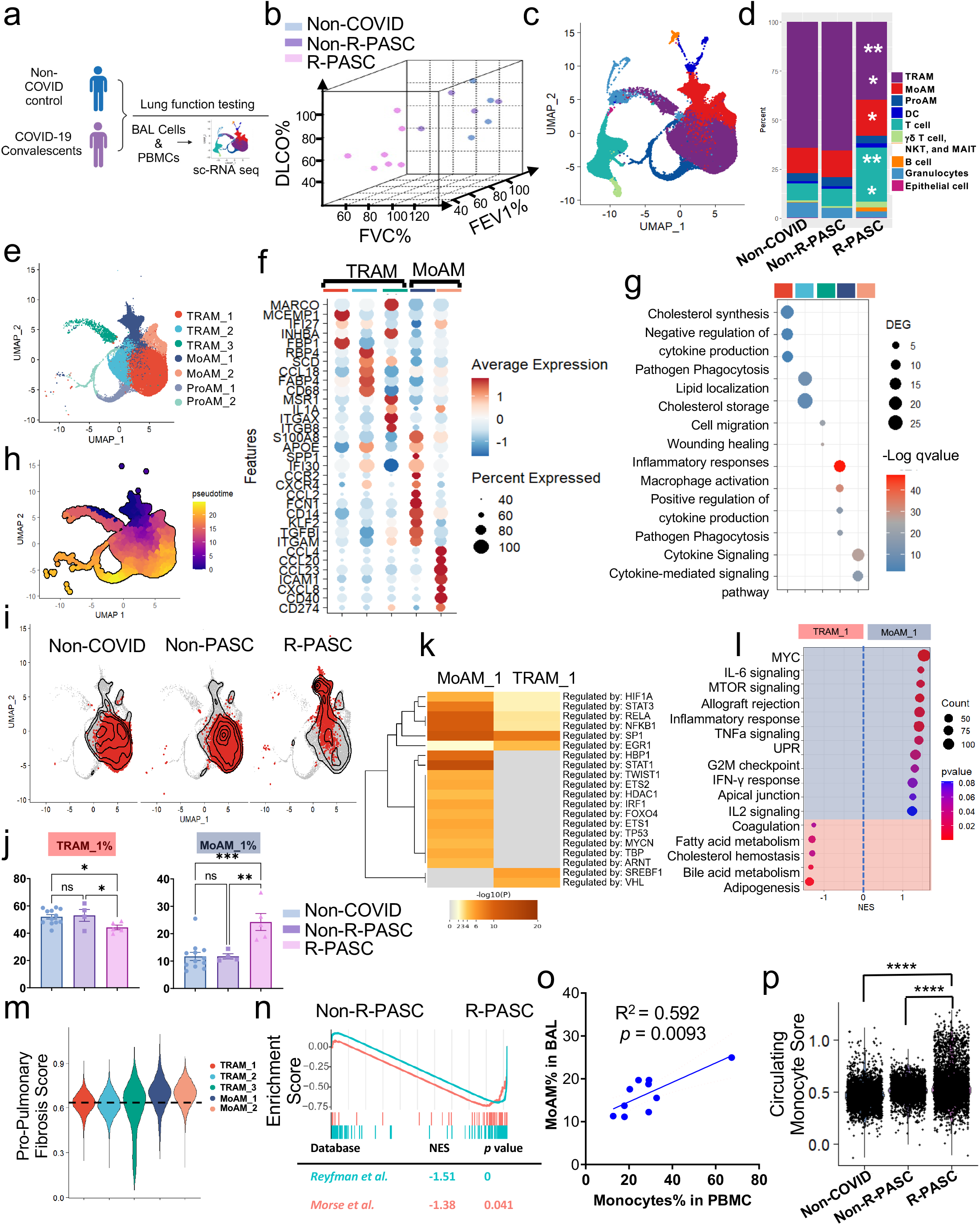
Accumulation of pro-inflammatory and pro-fibrotic MoAM in R-PASC. **a**, The schematic of the experimental procedure. **b**, Three-dimensional distributions of the lung function test results. DLCO, diffusing capacity of the lungs for carbon monoxide; FVC, Forced vital capacity; FEV1, Forced expiratory volume in the first second. **c**, UMAP plots showing integrated analysis of BAL cells grouped by the lung function, including four non-R-PASC, five R-PASC, two non-COVID control, and these data were integrated with published sc-RNA-seq dataset^17^ that contain 10 healthy BAL samples before the pandemic. **d**, Stacked bar plots showing the proportion of indicated cell types in the BAL among non-COVID, non-R-PASC and PASC groups. **e**, The UMAP plot of BAL macrophage populations. **f**, Signature gene expression by each subcluster of BAL macrophages. **g**, Enriched pathways in BAL macrophage clusters. **h**, Pseudotime analysis of BAL macrophage differentiation. **i**, Contour plots showing the density of cell clusters in the UMAP among non-COVID, non-R-PASC and PASC groups. **j**, Proportions of TRAM_1 and MoAM_1clusters in BAL macrophages among non-COVID, non-R-PASC and R-PASC groups. **k**, TRRUST predicted transcriptional regulation between TRAM_1 and MoAM_1cluster. **l**, Differential pathways enriched in MoAM_1cluster and TRAM_1 cluster. **m**, Relative score of Pro-Pulmonary Fibrotic Macrophage gene signature in BAL macrophage clusters. **n**, GSEA of Pulmonary Fibrosis-related macrophage gene sets^27, 28^ between non-R-PASC and R-PASC MoAM_1 cells. **o**, Correlation of BAL MoAM_1 proportions and PBMC monocytes percentages. **p**, PBMC monocyte feature assessment in MoAM_1 cells from indicated groups. Data are represented mean ± SEM unless otherwise indicated. Significance were tested by one-way ANOVA with Tukey’s adjustment or Wilcoxon test, ∗p < 0.05; ∗∗p < 0.01; ∗∗∗p < 0.001; ∗∗∗∗p < 0.0001.

Convalescents with either FEV1 or FVC values below 80% of predicted were categorized as R-PASC patients, while the rest were labeled as non-R-PASC (Fig. 1b and Table 1). To discern the underlying differences in the immune landscape between R-PASC and non-R-PASC convalescents, we conducted scRNAseq on bronchoalveolar lavage (BAL) and peripheral blood mononuclear cells (PBMC). In total 85,971 BAL cells and 101,296 PBMCs (from five R-PASC, four non-R-PASC, and two control individuals) were analyzed after integrating with published healthy datasets (additional ten individuals)^16, 17^. Ten major BAL cell clusters and ten PBMC clusters were identified (Fig. 1c, and Extended data Fig. 1a, b, c). In comparison to non-infected controls and non-R-PASC convalescents, R-PASC patients displayed an increased proportion of monocyte-derived alveolar macrophages (MoAM) and T cells in BAL, and monocytes in PBMC (Fig. 1 d, and Extended data Fig.1d, e, f). Conversely, tissue resident alveolar macrophage (TRAM) counts significantly diminished in the BAL of R-PASC patients (Fig. 1d and Extended data Fig. 1d).

Non-R-PASC BAL cells and PBMCs exhibited upregulated pathways such as IL2 signaling, KRAS signaling, glycolysis, and the unfolded protein response (UPR) compared to controls (Extended data Fig. 2a, b). BAL cells from R-PASC patients, however, predominantly highlighted pathways associated with cell proliferation and inflammation, like the G2M checkpoint, MTOR pathways, and allograft rejection (Extended data Fig. 2a). As the result, TRAM characteristic genes, such as *PPARG*, *FABP4*, and *MARCO*, were prominently expressed in non-R-PASC BAL cells. In contrast, genes indicative of cytotoxic T cells, like *NKG7*, *CCL5*, *GZMK*, and *CXCR6*, were prevalent in BAL cells from R-PASC patients (Extended data Fig. 2c, d), consistent with what we and others have reported that T cell signatures are enriched in R-PASC^15, 18^. GSEA analysis between R-PASC and non-R-PASC groups illustrated that pathway prominent in non-R-PASC BAL cells revolved around AM-driven tissue homeostasis, while R-PASC cell pathways from both BAL and peripheral blood leaned towards tissue reactivity and inflammation (Extended data Fig. 2e, f). Notably, the abundance of BAL MoAM and PBMC monocytes, coupled with the decreased TRAM count, correlated with compromised lung function in R-PASC patients, including a reduction in FEV1%, FVC%, and DLCO% (Extended data Fig. 2g, h). Taken together, these data suggest that R-PASC is characterized by marked alteration of immune cell composition and inflammatory responses in the respiratory tract compared to those of non-infected controls or infected subjects without R-PASC.

**Figure 2.**
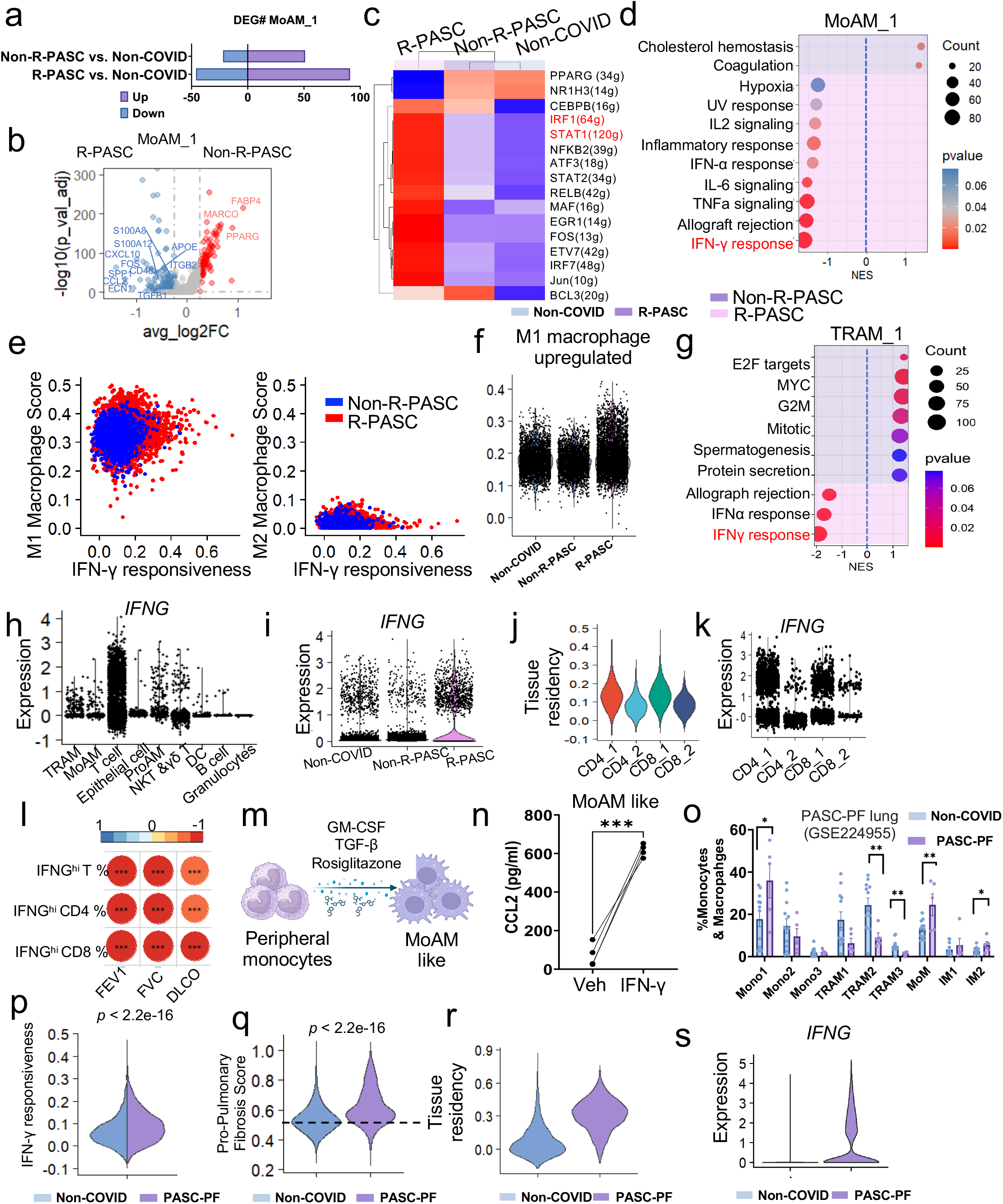
T cell-derived IFN-γ promotes MoAM recruitment, phenotype and polarization in R-PASC. **a**, Differentially expressed gene (DEG) counts in MoAM_1 cells from indicated comparison. **b**, Volcano plot showing the differentially expressed genes between non-R-PASC and R-PASC MoAM_1 cells. **c**, SCINEC analysis of transcriptional regulation of MoAM_1 from indicated groups. **d**, Dot plot showing enriched pathways in non-R-PASC and R-PASC MoAM_1 cells. **e**, Scatter plots showing IFN-γ responsiveness score and M1 (left) or M2 (right) macrophage features in MoAM_1 cells from indicated groups. **f**, M1 macrophage feature score of non-COVID, no-R-PASC and R-PASC MoAM_1 cells. **g**, Gene sets enriched in non-R-PASC and R-PASC TRAM_1 cells. **h**, *IFNG* expression in indicated cell types from BAL. **i,** Violin plot showing *IFNG* expression in BAL T cells from indicated groups. **j**, **k,** Tissue residency score (**j**) and *IFNG* expression (**k**) of BAL CD4^+^ conventional and CD8^+^ T cell subclusters. **l,** Correlation of IFNG highly expressed cell proportion with lung functional parameters. **m,** Schematic of peripheral monocyte derived MoAM-like cells treated with recombinant IFN-γ. **n**, CCL2 concentration in the culture medium from IFN-γ or Vehicle treated MoAM like cells. **o,** Proportion of indicated macrophage clusters in indicated lung tissue from non-COVID or PASC patients with pulmonary fibrosis (PASC-PF) in GSE224955. **p,** IFN-γ responsiveness score of MoAMs from Non-COVID or PASC-PF patient lung tissues in GSE224955. **q,** Relative score of pro-pulmonary fibrotic macrophage gene signature in MoAM cells in indicated groups in GSE224955^4, 41, 42^. **r, s,** Tissue residency score (**r**) and *IFNG* expression (**s**) in T cells from indicated lung tissues in GSE224955. Data are represented mean ± SEM. Significance were tested by paired *t* test, Wilcoxon test, or one-way ANOVA with Tukey’s adjustment, * p < 0.05, ** p < 0.01, *** p < 0.001, and **** p < 0.0001.

### Pro-fibrotic monocyte-derived macrophages accumulate in R-PASC

We previously have reported scRNAseq analysis of purified T cells in this cohort; here, we observed BAL T cells from R-PASC patients showed terminal differentiated features with upregulating *KLRG1*, *GZMK* expression, and B cells from R-PASC BAL showed noticeable differentially expressed genes (DEGs) compared with non-R-PASC counterparts (Extended data Fig. 2i, j). Subsequent analysis was focused on macrophages, as those cells have been suggested to link with the development of tissue fibrosis in animal models^19, 20^. Functional correlation analysis revealed a contrasting distribution of macrophage populations in convalescents with or without R-PASC. In order to gain a higher resolution of BAL macrophages in R-PASC, we further subclustered macrophages into seven subclusters with distinct gene expression profiles including three TRAM clusters (TRAM_1, TRAM_2, TRAM_3), two proliferating AM clusters (ProAM_1, ProAM_2) and two monocyte-derived macrophage populations (MoAM_1 and MoAM_2) (Fig. 1e). Among two monocyte derived macrophage clusters, MoAM_1 expressed high levels of *APOE*, *CD14* and *FCN1*, indicative of a transitory differentiation state from monocytes to macrophages. This cell cluster was marked by a high expression of alarmins (*S100A8*), inflammatory chemokines (*CCL2*) and chemokine receptors (*CCR2*, *CXCR4*) (Fig. 1f). Additionally, MoAM_1 were associated with inflammatory responses, macrophage activation, cytokine production, and pathogen phagocytosis (Fig. 1g). Notably, MoAM_1 population expressed high levels of *SPP1*, which encodes Osteopontin—a multifunctional matricellular protein and cytokine seen in macrophages across diverse pathologies, implicated as a pivotal factor in tissue damage and fibrosis^21, 22, 23^. The MoAM_2 cluster showed enhanced expression of *CD274* (PD-L1), *CD40*, and *ICAM1*, primarily linking to cytokine-mediated signaling (Fig. 1f, g). We also identified three tissue resident alveolar macrophage types (TRAM_1, TRAM_2, and TRAM_3), characterized by elevated expression of *FBP1*, *FABP4*, *CD68* and *MARCO* (Fig. 1f)—traits characteristic of TRAMs^24^. These cells were predominantly enriched with cholesterol synthesis regulation, wound healing and lipid metabolism pathways (Fig. 1g). Furthermore, two proliferative AM (ProAM) populations were characterized by the expression of cell-cycle-related genes (*MKI67*, *NUSAP1* and *CDK1*).

Trajectory analysis posited that MoAM_1 likely differentiated towards TRAM_1 (Fig. 1h). When compared to control or non-R-PASC donors, BAL cells from R-PASC patients showed a marked rise in the MoAM_1 population and a decreased TRAM_1 population (Fig. 1i, j and Extended data Fig. 3a). Furthermore, there was a negative association between the enrichment of MoAM_1 and lung function recovery (Extended data Fig. 3b). Transcription regulation analysis^25^ elucidated that, relative to TRAM_1, DEGs of MoAM_1 are modulated by hypoxia (mediated by transcription factors (TFs) like HIF1A, VHL), and proinflammatory cytokine signaling (TFs such as STAT3, STAT1, IRF1 and NFKB1) (Fig.1k). This is in agreement with GSEA, which indicated enrichment in IFN-γ, IL-6, and TNF signaling within the MoAM_1 cluster (Fig. 1l).

**Figure 3.**
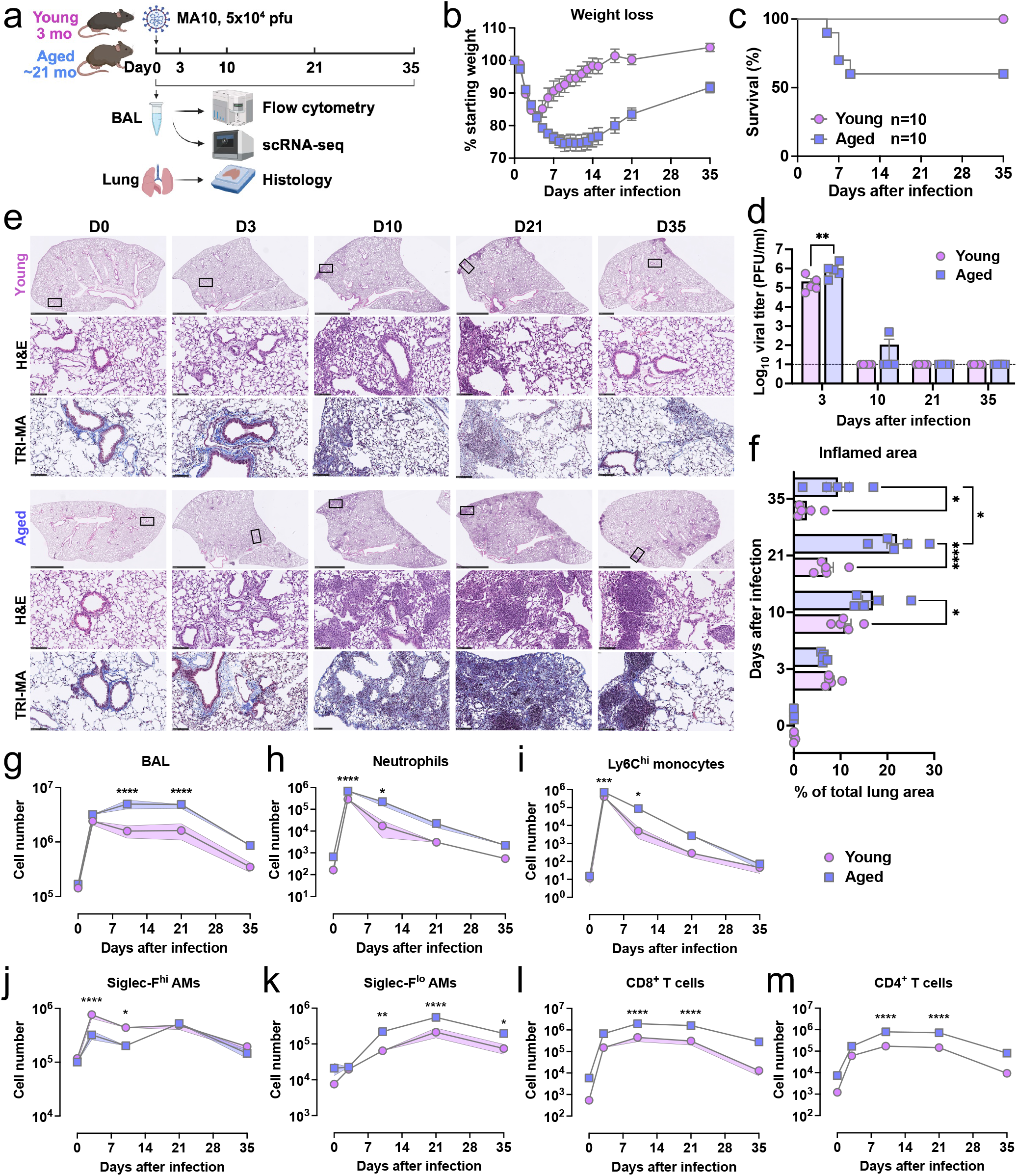
A model of postacute SARS-CoV-2 pulmonary sequelae in aged C57BL/6 mice. **a**, Schematic overview of SARS-CoV-2 MA10 infection experimental design. **b, c**, Weight loss (**b**) and survival (**c**) of young and aged mice were monitored. **d**, Viral titers in the respiratory track (BAL) were determined by plaque assay on Vero E6 cells. Symbols represent individual mice. The dashed line indicates the detection limit. **e**, Representative images of histopathology are shown. H&E indicates hematoxylin and eosin staining. Masson’s trichrome (TRI-MA) staining highlights fibrotic collagen deposition. The scale bar indicates 2.5 mm for the whole lung lobe or 100 mm for the zoomed area. **f**, QuPath quantification of inflamed area in the lungs for indicated time points as in **e**, represented as the percentage of total lung area. **g - m**, Total BAL cell counts (**g**), neutrophils (**h**), Ly6C^hi^ monocytes (**i**), Siglec-F^hi^ AMs (**j**), Siglec-F^lo^ AMs (**k**), CD8^+^ T cells (**l**), and CD4^+^ T cells (**m**) in BAL of MA10-infected young and aged mice at the indicated time points. Data are represented mean ± SEM unless otherwise indicated. Significance were tested by two-way ANOVA with Tukey’s adjustment for multiple comparisons, * p < 0.05, ** p < 0.01, *** p < 0.001, and **** p < 0.0001.

It was indicated that MoAM generally adopts a pro-fibrotic phenotype during severe COVID-19 acute respiratory distress syndrome (ARDS)^26^; however, it remains uncertain if the accumulated MoAM cells in R-PASC patients retain these pro-fibrotic characteristics after the resolution of ARDS. We thus evaluated pro-pulmonary fibrosis macrophage core genes in BAL macrophages ^27, 28, 29^. MoAM_1 consistently scored the highest among the five macrophage populations examined (Fig.1m, and Extended data Fig. 3c, d). Additionally, the R-PASC patients derived MoAM_1 subset displayed notable enrichment of pro-pulmonary fibrosis macrophage gene sets (Fig. 1n, and Extended data Fig. 3e).

The heightened expression of the monocyte chemoattractant *CCL2* in MoAM_1 cells hints at a feedback mechanism for monocyte recruitment and subsequent macrophage differentiation (Fig. 1f, and Extended data Fig. 3f-h). The positive correlation between BAL MoAMs and circulating monocytes further underscores this hypothesis (Fig. 1o). However, no significant correlation was observed between circulating T cells, NK cells, B cells, and their respiratory equivalents (Extended data Fig. 3i). To further explore the recruitment of MoAMs from circulation, transcriptional similarity was assessed between circulating monocytes and BAL macrophages, this assessment demonstrates that the MoAM_1 subset in R-PASC patients bears a strong resemblance to circulating monocytes (Extended data Fig. 3j, and Fig. 1p). Collectively, our comprehensive single-cell transcriptome analysis revealed that post-COVID-19 lung sequelae are associated with marked dysregulation in the pulmonary macrophage population, with R-PASC individuals exhibiting increased presence of proinflammatory and pro-fibrotic MoAMs, likely due to persistent recruitment of monocytes to the respiratory tract and/or incomplete differentiation of these monocytes to gain the mature TRAM phenotype. Notably, a recent pre-print manuscript with a larger cohort of BAL donors also identified that increased pulmonary MoAMs were associated with persistent respiratory symptoms and radiographic abnormalities ^30^.

### Resident T cell-derived IFN-γ promotes MoAM recruitment and phenotype in R-PASC

Differential analysis between BAL macrophages from R-PASC and non-R-PASC individuals elucidated a marked enrichment of inflammatory responses and cytokine signaling in R-PASC-derived macrophages. In contrast, lipid metabolic pathways were predominantly evident in macrophages from non-R-PASC donors (Extended data Fig. 4a). Focusing on MoAM_1 subset, there were more DEG counts between R-PASC to non-COVID groups than those gene counts between non-R-PASC and non-COVID groups (Fig. 2a). Direct comparison of MoAM_1 cells from R-PASC and non-R-PASC groups showed an upregulation of TRAM associated genes, namely *FABP4*, *PPARG*, and *MARCO* in MoAM_1 cells from non-R-PASC donors. In contrast, MoAM_1 cells from R-PASC patients exhibited increased expression of inflammatory chemokines (*CCL2*, *CXCL10*), inflammatory regulators (*FCN1*, *S100A12*, *S100A8*), the lung fibrosis factor *SPP1*, and the monocyte-derived alveolar macrophage marker *APOE* (Fig. 2b). These data suggest that MoAM_1 from R-PASC groups may be more arrested toward the TRAM differentiation. We then applied SCENIC^31^ to discover potential transcriptional regulators that modulate the differential gene expression in MoAM cells from the three groups. We found that MoAM_1 cells in R-PASC group showed increased STAT1, IRF1, IRF7, and NFKB2 TF activities (Fig. 2c), consistent with enriched proinflammatory gene sets (Fig. 2d). Notably, a prominent trait of R-PASC MoAM_1 cells was the heightened IFN-γ response among enriched pathways (Fig. 2d). *IFNGR2*—a critical determinant for IFN-γ responsiveness^32, 33^ —was elevated, further indicating an upregulated IFN-γ response in R-PASC patient-derived MoAM_1 cells (Extended data Fig. 4b, c).

**Figure 4.**
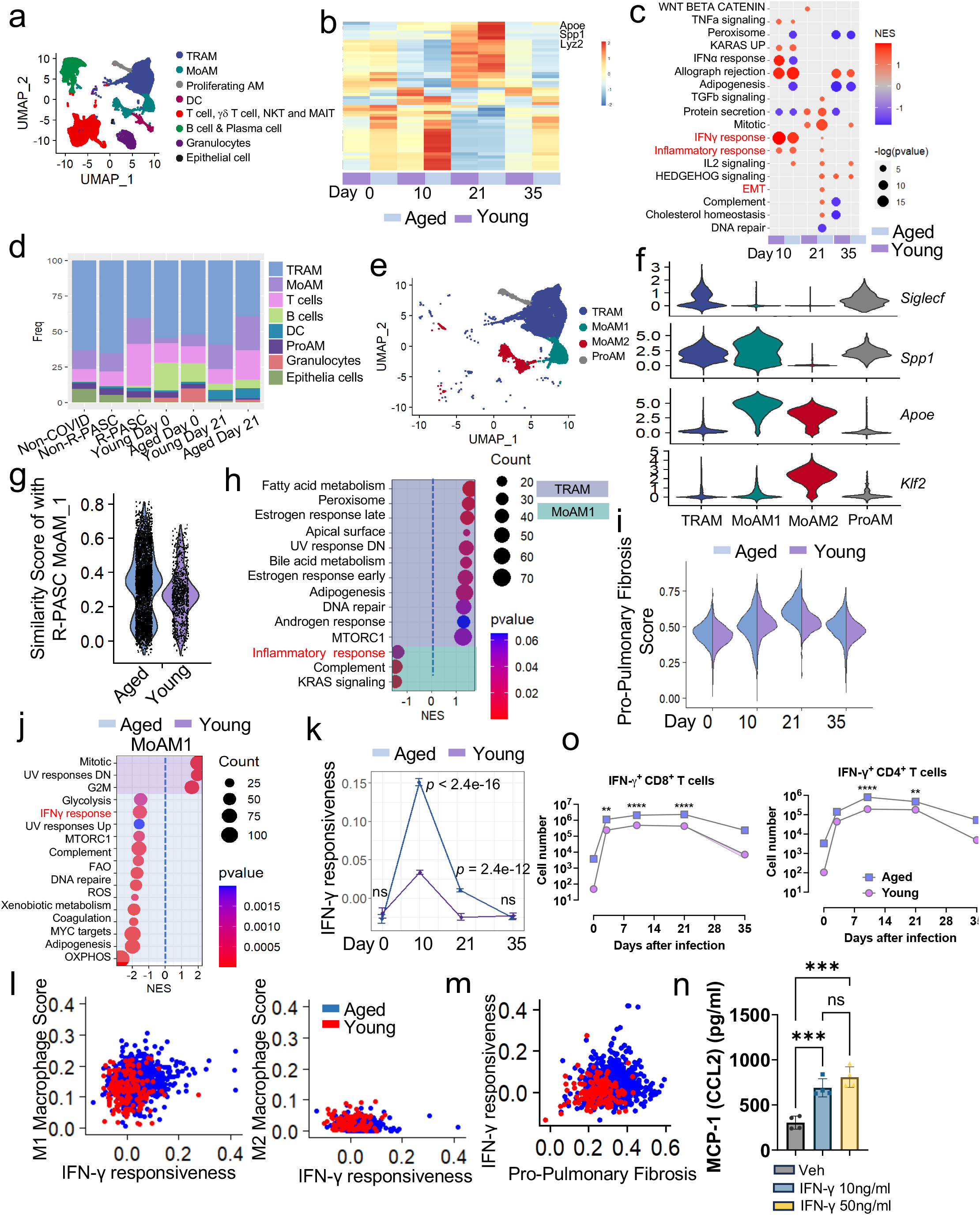
Comparative sc-RNA-seq analysis of human and mouse R-PASC BAL cells. **a,** The integrated UMAP of BAL cells from young and aged C57BL/6J mice at 0, 10, 21 and 35 dpi. **b,** Heatmap showing the top 50 variable genes of BAL cells in young or aged mice at indicated time points. **c,** The enriched pathways in BAL cells at 10, 21 and 35 dpi compared with uninfected (day 0). **d,** Stacked bar plots showing the proportion of indicated cell types in BAL of human R-PASC, non-R-PASC and infected mice. **e,** The integrated UMAP of BAL macrophages from young and aged C57BL/6J mice at 0, 10, 21 and 35 days post MA10 infection. **f,** Violin plot of indicated gene expression in the BAL macrophage subclusters. **g,** The similarity score of human MoAM_1 features in young or aged C57BL/6J mouse MoAM1 cells at 21 dpi. **h,** Gene set enrichment in MoAM1 and TRAM clusters at 21 dpi. **i,** Relative score of pro-pulmonary fibrotic macrophage gene signature in MoAM1 cells at indicated group and time points. **j,** Enriched gene sets of MoAM1 clusters from young or aged C57BL/6J mice at 21 dpi. **k,** INF-γ responsiveness score of MoAM1 from young or aged C57BL/6J mice at indicated time points post MA10 infection. **l,** Scatter plots showing IFN-γ responsiveness score and M1 (left) or M2 (right) macrophage features in MoAM1 cells from young or aged C57BL/6J mice at 21 dpi. **m,** Scatter plot showing IFN-γ responsiveness and pro-pulmonary fibrosis scores in MoAM1 cells from young or aged C57BL/6J mice at 21 dpi. **n,** Culture medium MCP-1 (CCL2) concentration at 24 hours post bone marrow derived MoAM-like cells exposure to IFN-γ. **o,** Total cell counts of IFN-γ-producing CD8^+^ T and CD4^+^ T cells in young or aged mice infected with MA10 at indicated time points. Data represent the mean ± SEM. Data were analyzed by two-way ANOVA or unpaired *t* test, * p < 0.05, ** p < 0.01, and **** p < 0.0001.

Additionally, IFN-γ responsive MoAM_1 cells demonstrated an elevated propensity for pro-pulmonary fibrosis in R-PASC patients (Extended data Fig. 4d). Macrophage polarization, influenced by factors such as IFN-γ, and IL-4, gives rise to distinct proinflammatory (M1) or pro-fibrotic (M2) gene expression profiles^34, 35^. Generally, compared with M1 polarized macrophages, M2 macrophages and Th2-driven responses are critical in the areas of lung fibrosis^36, 37, 38, 39^. We found that MoAM_1 cells exhibited the upregulated M1 differentiation features while gaining the pro-fibrotic profile and IFN-γ responsiveness. Conversely, the M2 features of MoAM_1 were low (Fig. 2e, and Extended data Fig. 4e). Furthermore, the MoAM_1 cells from R-PASC patients have the highest M1 score, and a comparable M2 score comparing the donors without COVID-19 or R-PASC (Fig. 2f, and Extended data Fig. 4f). Together, these data indicate that R-PASC MoAM_1 cells are M1-polarized. Of note, similar IFN-γ response enrichment was discerned in other lung and circulatory cell types, including TRAM_1, respiratory epithelial cells, MoAM_2, and circulating monocytes (Fig. 2g, and Extended data Fig. 4g-i). These data revealed a widespread IFN-γ response across many respiratory cell types in R-PASC individuals, potentially due to the presence of an IFN-γ abundant milieu in the respiratory tract of those individuals.

To explore the cellular source of IFN-γ, *IFNG* expression levels were assessed in BAL cells, T cells were revealed as the major cell type harboring high levels of *IFNG* transcripts (Fig. 2h). Correlating with the increased prevalence of CD4^+^ conventional T cells and CD8^+^ T cells in R-PASC BAL (Extended data Fig. 4j, k), there was an augmented *IFNG* expression from CD4^+^ conventional and CD8^+^ T cells (Fig. 2i and Extended data Fig. 4l). Remarkably, most of these *IFNG-*expressing T cell subsets exhibited tissue-resident characteristics (Extended data Fig. 4m and Fig. 2j, k). Consequently, compared to those of the non-R-PASC group, there appeared to be elevated IFN-γ concentrations in BAL from R-PASC patients, and R-PASC BAL-derived CD4^+^ and CD8^+^ T cells showcased enhanced IFN-γ protein production upon antigenic stimulation (Extended data Fig. 4n, o), although the limited sample size here prevents us to draw a firm conclusion. Additionally, correlation analysis in the scRNAseq data revealed that BAL *IFNG*-expressing T cell proportions showed a negative association with lung function recovery (Fig. 2l). The expression of type 2 and type 3 related cytokines, which were associated with M2 macrophage polarization, was almost undetectable among three groups of donors (Extended data Fig. 4p), further indicating that the IFN-γ mediated type 1 immune responses likely promote lung fibrotic responses in R-PASC.

MoAM cells from R-PASC patients exhibited overexpressed *CCL2* (Extended data Fig. 3h and Fig. 2b), indicating continuous monocyte recruitment. To determine if IFN-γ plays a role in this process, we differentiated MoAM like cells from human peripheral monocytes after the treatment TGF-β, GM-CSF, and PPARG agonist Rosiglitazone *in vitro*^40^ (Fig. 2m). When compared to monocytes, *in vitro* differentiated MoAM like cells displayed increased expression of AM cell markers, such as CD169 and CD68 (Extended data Fig. 4q). After exposed to recombinant human IFN-γ, MoAM-like cells exhibited a significant rise in CCL2 expression compared to untreated cells (Fig. 2n and Extended data Fig. 4r). These data suggest that IFN-γ abundant microenvironment amplifies CCL2 production by MoAMs, further boosting or sustaining monocyte recruitment to the respiratory tract in R-PASC. In summation, our data reveal that individuals with R-PASC are characterized with a potential exuberant communication between respiratory resident T cells and myeloid cells (particularly MoAM cells) mediated by IFN-γ, driving persistent monocyte recruitment and subsequent M1-like polarization in monocyte-derived macrophages.

To further explore the altered immune status in lung tissues from R-PASC patients, we analyzed a lung scRNAseq dataset from a cohort of five patients with extensive lung fibrosis (PASC-PF) that requires lung transplantation^4^ (Extended data Fig. 5a). Analysis of lung monocytes and macrophages revealed substantial alterations in cellular composition in PASC-PF lungs compared to those non-COVID controls^41, 42^. We observed markedly increased monocyte subclusters Mono1 and MoAM and diminished TRAM2 and TRAM3 clusters in PASC-PF lungs (Extended data Fig. 5b, and Fig. 2o). This shift mirrors the macrophage composition observed in BAL samples from our cohort. Notably, MoAM from PASC-PF lungs displayed increased IFN-γ responsiveness, enhanced pro-fibrotic characteristics and a bias towards M1 differentiation (Fig. 2 p,q, and Extended data Fig. 5c). Further assessment also pinpointed that T cells with increased tissue-resident characteristics were the major cellular source of *IFNG* expression in the lung tissues during PASC-PF (Fig.2 r, s and Extended data Fig.5d). Thus, exuberant IFN-γ-mediated pulmonary T-macrophage communications appear to be present in two cohorts of respiratory PASC cohorts.

**Figure 5.**
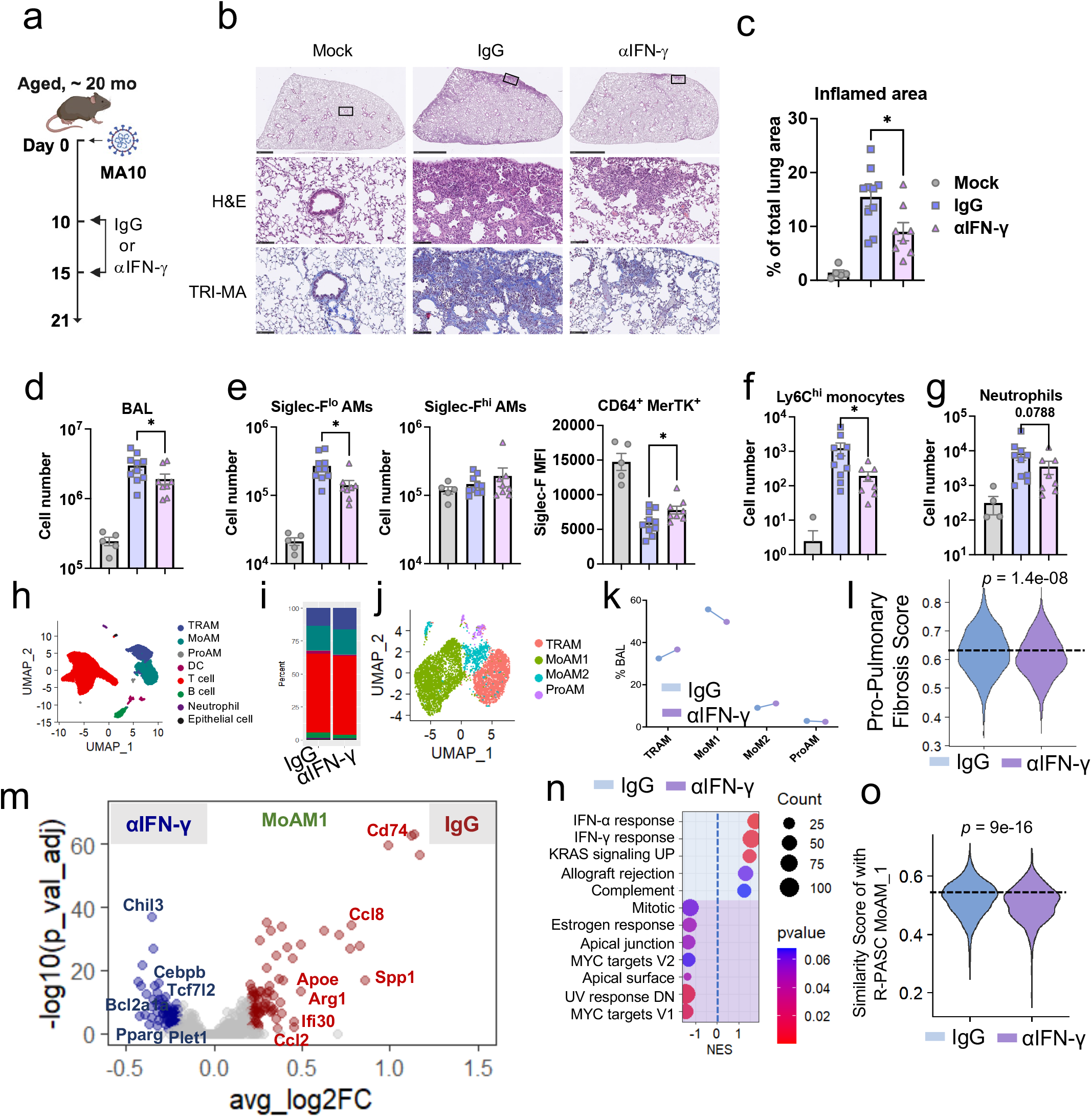
Therapeutic targeting persistent IFN-γ mitigates pulmonary pathology postacute SARS-CoV-2 infection. **a,** Experimental setup for evaluating the role of INF-γ in MA10 infection-induced lung pathology in aged B6 mice. **b,** Representative images of lungs stained for H&E and trichrome from aged mice treated with IgG or αIFN-γ. Scale bars, 2.5 mm for the whole lung lobe, and 100 mm for the zoomed area. **c,** Quantification of inflamed lung area from IgG- or αIFN-γ-treated aged mice. Mock, n = 5 mice; IgG, n = 10 mice; αIFN-γ, n = 8 mice, pooled from two independent experiments. **d - g,** Total BAL cell counts (**d**), Siglec-F^lo^ AMs and Siglec-F^hi^ AMs (**e**), Ly6C^hi^ monocytes (**f**), and neutrophils (**g**) in BAL of MA10-infected aged mice treated with IgG or αIFN-γ. **h**, The UMAP of BAL cells from IgG or αIFN-γ treated aged mice. **i**, Stacked bar plots showing the proportion of indicated cell types in BAL cells from IgG or αIFN-γ treated aged mice. **j**, The UMAP of BAL macrophages from IgG or αIFN-γ treated aged mice. **k**, The proportion of indicated macrophage clusters in BAL cells from IgG or αIFN-γ treated aged mice. **l,** Relative score of pro-pulmonary fibrotic macrophage gene signature in MoAM1 cells at indicated group post treatment. **m,** Volcano plot showing the differentially expressed genes of MoAM1 cells from IgG or αIFN-γ treated aged mice. **n,** Enriched gene sets of MoAM1 clusters from IgG or αIFN-γ treated aged mice. **o,** The similarity score of human MoAM_1 features in IgG or αIFN-γ treated aged mouse MoAM1 cells. Data represent the mean ± SEM. Data were analyzed by onw-way ANOVA, unpaired *t* test, or Wilcoxon test, * p < 0.05, ** p < 0.01, and **** p < 0.0001.

### Aged C57BL/6J mice manifest pulmonary sequelae after acute SARS-CoV-2 infection

To gain insight into the underlying mechanisms, we leveraged a mouse adapted strain of SARS-CoV-2, MA10, generated from the ancestral Wuhan isolate^43^. In humans, advanced age is a known risk factor for developing severe and lethal disease following SARS-CoV-2 infection^44^. To determine this age dependent susceptibility to SARS-CoV-2 infection, we infected both young (3-mo-old) and aged (21-mo-old) female mice with 5×10^4^ pfu MA10 virus and monitored their weight changes (Fig. 3a). Infection of young mice resulted in an approximately 15% weight loss followed by rapid recovery, whereas aged animals experienced more substantial and prolonged weight loss (Fig. 3b). Furthermore, none of the young mice succumbed to the infection, while 40% of the aged mice succumbed to the infection or reached the humane euthanasia criteria (Fig. 3c).

To elucidate the age-associated disease progression, we collected tissues at various time points to capture both the acute phase and post-acute phase of infection, and assessed lung pathology and immune responses in the respiratory tract (Fig. 3a). Three days post infection (dpi), virus replication in the lungs of aged mice increased about 5-fold compared to young mice and there was no detectable infectious virus by 10 dpi in all mice except for one aged mouse (Fig. 3d). Additionally, histological examination revealed that young mice exhibited subpleural lesions from days 10 to 35. The lung pathology in young mice peaked at 10 dpi and essentially all recovered at 35 dpi. In contrast, aged mice displayed more severe lung pathology with considerable lung inflammation at 21 dpi when the virus had been completely cleared after primary infection. Lung pathology in aged mice remained noticeable at 35 dpi, although largely resolved from day 21 (Fig. 3e, f). Similarly, heightened collagen deposition was also evident in the aged mice at 21 dpi (Fig. 3e). These data suggest that aged C57BL/6J mice manifest pulmonary inflammatory and fibrotic sequelae after acute SARS-CoV-2 infection.

We next compared immune cell recruitment in the BAL of young and aged mice. MA10 infection triggered a rapid infiltrating of cells into the respiratory tract in both age groups at the evaluated infection time points. However, aged mice exhibited a notably higher cell count upon 3 dpi (Fig. 3g). In addition, a significantly elevated number of neutrophils and inflammatory monocytes were seen in infected aged mice compared with young counterparts (Fig. 3h, i, and Extended data Fig. 6a, b). Previous studies have shown that influenza virus infection can lead to a reduction in Siglec-F^hi^ TRAMs, subsequently instigating the emergence of Siglec-F^lo^ MoAMs^45, 46^. Indeed, our observations align with these findings, demonstrating a significant reduction in the percentage of TRAMs and a marked increase in MoAMs in both age groups. Notably, aged mice showed a more pronounced deficit in TRAMs and a greater prevalence of MoAMs in comparison to their young counterparts (Fig. 3j, k, and Extended data Fig. 6a, b), suggesting a more proinflammatory response in aged animals. Moreover, we observed significantly higher CD8^+^ and CD4^+^ T cells in aged mice. Intriguingly, while their proportion is largely reduced compared to young mice, overall spike-specific CD8^+^ T cell counts are higher in aged mice, indicating a more prominent bystander CD8^+^ T cell response in aged mice (Fig. 3l, m, and Extended data Fig. 6c, d). Together, these data indicate that the post SARS-CoV-2 pulmonary sequelae model developed in aged C57BL/6 mice is characterized by similar respiratory proinflammatory cell profiles as those R-PASC individuals.

**Figure 6.**
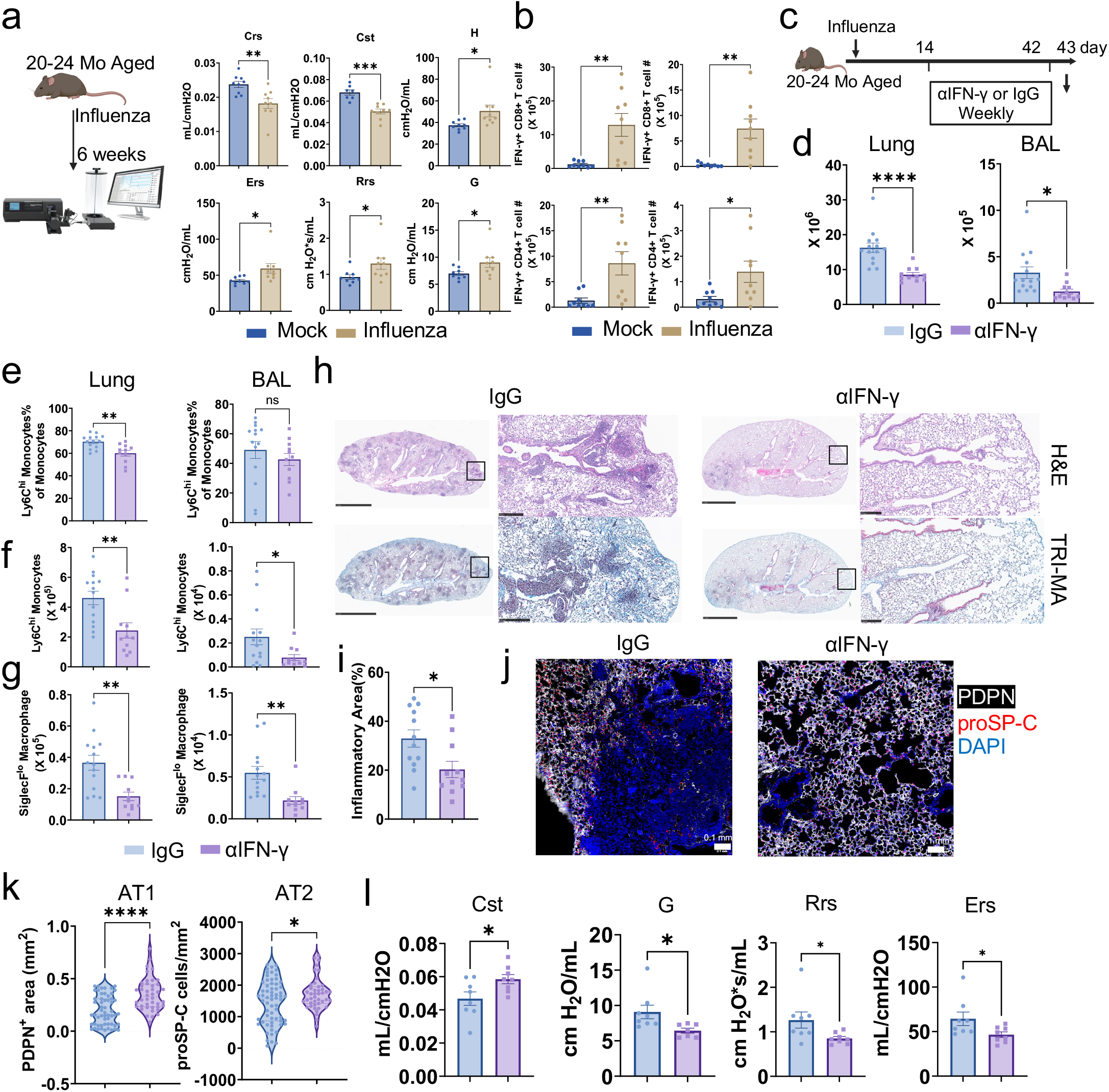
IFN-γ blockade promotes lung functional recovery in a persistent sequelae model caused by influenza pneumonia. **a,** Evaluation of respiratory compliance (Crs, Cst), tissue resistance (H), respiratory Elastance (Ers), respiratory Resistance (Rrs) and respiratory damping (G) in aged naïve or 6 weeks post infected C57BL/6J mice with flexiVent, n= 9, pooled from two independent experiments. **b,** IFN-γ producing lung-resident (left) and BAL (right) CD8^+^ and CD4^+^ T cell counts in aged naïve or 6 weeks post influenza infected C57BL/6J mice. n= 9, pooled from two independent experiments. **c,** Schematics for evaluating the role of IFN-γ in influenza infection-induced lung pathology in aged C57BL/6J mice. **d,** Total lung and BAL cell counts in infected aged mice post treatment, n= 11-14, pooled from three independent experiments. **e** and **f,** Proportion (**e**) and counts (**f**) of Ly6c^hi^ monocytes in the lung and BAL post treatment. **g,** Siglec-F^lo^ AMs counts in the lung and BAL post treatment, n= 11-14, pooled from three independent experiments. **h,** Representative images of lungs stained for H&E and trichrome from aged mice treated with IgG or αIFN-γ. Scale bars, 2.5 mm for the whole lung lobe, and 250 mm for the zoomed area. **i**, Quantification of inflamed lung area from IgG- or anti-IFN-γ-treated aged mice. **j,** Immune fluorescent staining of PDPN and proSP-C in the aged mouse lung sections post treatment. **k**, Quantification of PDPN staining positive aera proportion per mm^2^ (left) and proSP-C staining positive cell counts per mm^2^ (right) as in **j,** each dot represents one randomly picked field, n= 7, pooled from two independent experiments. **l**, Evaluation of compliance (Cst), damping (G), Resistance (Rrs) and Elastance (Ers) of the respiratory system in infected aged mice post treatment with flexiVent, n= 8, pooled from two independent experiments. Data represent the mean ± SEM. Data were analyzed by unpaired *t* test, * p < 0.05, ** p < 0.01, *** p < 0.001, and **** p < 0.0001.

### scRNAseq analysis of respiratory immune cells in mouse R-PASC

We next sought to understand the cellular and molecular profiles associated with this mouse R-PASC. To this end, sc-RNA-seq was performed on BAL cells from both young and aged C57BL/6J mice at various time-points post-infection (0, 10, 21, and 35 dpi). Subsequent analysis identified 8 distinct cell populations in the BAL post SARS-CoV-2 infection (Fig. 4a and Extended data Fig. 7a). Kinetic analysis revealed that T cell proportion showed an increase following infection in the aged mice. Moreover, an accumulation of MoAM populations, coupled with a decrease in TRAM populations in aged mice at 21dpi (Extended data Fig. 7b). MoAM cells were greatly diminished in the respiratory tract at 35 dpi, coinciding with the decreased lung pathology at 35 dpi.

Notably, differential gene expression profiles between young and aged mice were pronounced at 21 dpi (Fig. 4b). Genes previously identified as upregulated in the MoAM population from R-PASC patients, including *Apoe*, *Spp1*, and *Lyz2*, were markedly upregulated in the aged mice at 21 dpi, and subsided by 35 dpi—aligning with the observed recovery of lung pathology (Fig. 4b). Associated with these findings was persistent inflammation in BAL cells from aged mice at 21 dpi (Fig. 4c). This temporal pattern correlates with lingering inflammatory responses observed in BAL cells from aged mice at 21 dpi, when pathways like TGF-β response and Epithelial–mesenchymal transition (EMT) were uniquely enriched, potentially hinting at inflammation-associated fibrosis development^47^ (Fig. 4c). Additionally, the patterns of BAL cell type composition in the aged mice at 21 dpi exhibited a similarity with those of R-PASC individuals (Fig. 4d), reinforcing the usefulness of the model to investigate the immune mechanisms underlying R-PASC.

To delineate the altered macrophage dynamics during the progression of lung sequelae, we stratified macrophages into four distinct clusters: a TRAM cluster, two MoAM clusters (MoAM1 and MoAM2), and a proliferating AM cluster (ProAM) based on their gene expression (Fig. 4e). TRAM cells predominantly express *Siglecf*, a hallmark marker of mature tissue-resident AMs. MoAM1 cells, characterized by elevated expression of *Spp1* and *Apoe*, manifested a pronounced accumulation in aged mice relative to their younger counterparts at 21 dpi (Fig. 4f, and Extended data Fig. 7c). In contrast, MoAM2 cells exhibited rapid lung infiltration during the acute infection phase, accompanied by upregulated expression of the migratory regulator, *Klf2* (Fig. 4f, and Extended data Fig. 7c). Pathway enrichment analysis further highlighted the superior endocytic potential of TRAM cells, the enhanced proinflammatory signature of MoAM1 cells, and the heightened migration and cellular extravasation capabilities of MoAM2 cells (Extended data. Fig. 7d).

Subsequent assessment of transcriptomic similarities between MoAM cells in R-PASC and analogous MoAM1 from SARS-CoV-2 MA10 infected mice indicated a significant transcriptomic overlap, particularly in MoAM1 cells from aged mice at 21 dpi (Fig. 4g, and Extended data Fig. 7e). A comparative analysis between MoAM1 and TRAM cells from aged mice unveiled the inflammatory polarization of MoAM1 cells at 21 dpi (Extended data Fig. 7f and Fig. 4h). Moreover, MoAM1 cells showcased elevated chemotactic properties, aligning with increased *Ccl2* expression (Extended data Fig. 7g, h), indicating they might be differentiated from consistently recruited monocytes. A defining characteristic of human R-PASC MoAM_1 is its elevated pro-pulmonary fibrosis access score (Fig. 1m). We observed a similar pattern wherein MoAM1 cells from aged mice possessed the highest pro-fibrosis score across all assessed time points and cell types at 21 dpi (Fig. 4i and Extended data Fig. 7i, j).

Consistent with human scRNAseq analysis, GSEA of mouse BAL cells identified a sustained IFN-γ response across diverse cell types (BAL Macrophages, MoAM1, TRAM, and epithelial cells) in aged mice at 21 dpi (Fig. 4j, and Extended data Fig. 7k-m). However, this chronic IFN-γ signaling waned by week 5 (Fig. 4k), aligning with the timeline of lung pathology resolution (Fig. 3e). Similar to the human R-PASC MoAM_1 subset, IFN-γ responsiveness levels in MoAM1 cells positively correlated with M1 like macrophage differentiation and heightened pro-pulmonary fibrosis features in aged mice at 21 dpi (Fig. 4l, m), further suggesting the potential association of IFN-γ stimulation and post-viral fibrotic response. Coincident with human MoAM-like cells (Fig. 2n), there was an IFN-γ dependent Ccl2 production after bone marrow-derived MoAM-like cells were treated with IFN-γ *in vitro* (Fig. 4n, and Extended data Fig. 7n), indicating the role of IFN-γ in monocyte chemotaxis to the lung for the subsequent development into the proinflammatory and pro-fibrotic MoAM1 cells. Notably, similar to what was observed in the human scRNAseq analysis, T cells exhibiting tissue-resident memory (TRM) traits emerged as primary IFN-γ sources at 21 dpi (Fig. 4o, and Extended data Fig. 7o, p).

Taken together, our comparative scRNAseq analysis on respiratory samples from both COVID-19 convalescents and SARS-CoV-2 infected mice identified that, the immune landscape in aged mice three weeks after infection resembles the respiratory immune profile observed in R-PASC individuals, marked by skewed TRAM and MoAM composition, prolonged IFN-γ responses, and pro-pulmonary fibrosis attributes in MoAM cells.

### Therapeutic targeting of persistent IFN-γ mitigates post SARS-CoV-2 lung sequelae

Our comprehensive analysis, spanning clinical samples to SARS-CoV-2 MA10 infected mice, uncovers IFN-γ signaling as a key potential contributor to developing post-COVID-19 chronic lung complications. To directly test this, we treated the SARS-CoV-2 MA10 infected aged C57BL/6 mice with anti-IFN-γ intraperitoneally to block IFN-γ signaling after the clearance of primary viral infection (i.e. 10 dpi). We then measured pulmonary pathology and respiratory immune responses at 21 dpi, the time-point when pronounced pulmonary sequelae were observed after SARS-CoV-2 infection (Fig. 5a). The treatment ameliorated lung inflammatory and fibrotic pathology compared to IgG controls, indicating an overall improvement in outcomes (Fig. 5b, c). In line with this data, IFN-γ blockade resulted in a reduction in total BAL cell counts (Fig. 5d), attributed to a decrease in Siglec-F^lo^ MoAMs and inflammatory monocytes (Fig. 5e and f, and Extended data Fig. 8a), to a lesser impact on neutrophils (Fig. 5g, and Extended data Fig. 8a). There was no significant difference in the counts of total CD8^+^ T and CD4^+^ T cells (Extended data Fig. 8b). However, Spike-specific CD8^+^ T cells were significantly diminished after anti-IFN-γ treatment (Extended data Fig. 8c).

To interrogate alterations in the respiratory immune profile subsequent to IFN-γ signaling blockade, we performed sc-RNA-seq on BAL cells isolated from control or anti-IFN-γ treated mice at 21 dpi. Following IFN-γ neutralization, we noted an increase in the TRAM population (Fig. 5h, i, and Extended data Fig. 8d). To further explore the role of IFN-γ in MoAM recruitment and differentiation, macrophages were isolated and subsequently re-clustered into distinct clusters: one TRAM, two MoAMs, and one proliferative AM subcluster (Fig. 5j). Anti-IFN-γ administration resulted in an augmented TRAM population presence while diminished MoAM1 cells (Fig. 5k), mirroring observations from flow cytometric analyses (Fig. 5e, f). Pseudotime trajectories suggest a directional push by IFN-γ signaling blockade toward TRAM differentiation (Extended data Fig. 8e, f). Post anti-IFN-γ treatment, MoAM1 clusters markedly exhibited attenuated pro-pulmonary fibrotic characteristics (Fig. 5l, and Extended data Fig. 8g-i), characterized by suppressed expression of genes including *Arg1, Spp1, Ccl8 and Ccl2,* and elevated expression of genes favoring tissue homeostasis and repair, including *Pparg, Cebpb,* and *Plet1*^48^ (Fig. 5m). This gene modulation correlated with improved lung histopathology and diminished collagen deposition (Fig. 5b). Moreover, anti-IFN-γ treatment dampened inflammatory pathways within MoAM cells and curtailed typical MoAM1 features observed in R-PASC patients (Fig. 5n, o). In conclusion, direct targeting extended IFN-γ responses ameliorated pro-inflammatory and pro-fibrotic macrophage development, and dampened lung pathology in this murine model of post SARS-CoV-2 lung sequelae.

### IFN-γ is dispensable for post SARS-CoV-2 lung sequelae in BALB/c mice

Previously, it was shown that SARS-CoV-2 MA10 infection in aged BALB/c mice (1 year old) triggered potent respiratory inflammatory responses and lung pathology at 1 month post infection, which can persist up till 120 dpi^13^. Similar to the previous findings, we observed that SARS-CoV-2 MA10 infected aged BALB/c mice displayed pronounced chronic lung anomalies at 1 month post infection, including cell infiltration and collagen deposition in the lung (Extended data Fig. 9a-d). scRNAseq analysis of BAL cells revealed decreased TRAM cell proportion and accumulation of MoAM cells (Extended data Fig. 9e, f). However, pathway analyses did not reveal significant IFN-γ or proinflammatory responses in those MoAM cells even though persistent lung pathology was observed in this model (Extended data Fig. 9g-k). Additionally, MoAM1 cells in infected BALB/c mice did not show significant congruence with human MoAM_1 cells from R-PASC individuals (Extended data Fig. 9l), and pro-fibrotic attributes in these cells were comparable between infected and uninfected control groups (Extended data Fig. 9m). Furthermore, there was a lack of observable differences in *Ifng*-expression from T cells between infected and uninfected BALB/c mice (Extended data Fig. 9n). Finally, the blockade of IFN-γ signaling starting at 10 dpi *in vivo* did not ameliorate infection-induced chronic lung pathology in this model (Extended data Fig. 9o, p). Taken together, these results suggest that chronic lung conditions developed after SARS-CoV-2 infection in BALB/c mice might not be mediated by IFN-γ and/or IFN-γ responsive MoAMs, which is in sharp contrast to what were observed in R-PASC individuals or SARS-CoV-2 infected C57BL/6 mice.

### Inhibition of IFN-γ enhances lung functional recovery in a persistent sequelae model after influenza viral pneumonia

In contrast to the relatively transient lung inflammatory and fibrotic sequelae after SARS-CoV-2 infection in animal models, we previously found that influenza viral pneumonia can lead to more persistent chronic lung pathology (more than 60 dpi) in aged C57BL/6 mice, likely due to more extensive alveolar damage caused by mouse-adapted influenza (H1N1 A/PR8/34) virus than SARS-CoV-2 virus during the primary infection in mice ^49^ ^50^. In this model, there was evident impairment in lung compliance (Crs, Cst), increased tissue elastance (H, Ers) and respiratory resistance (Rrs), and heightened trend of respiratory damping (G) at 6 weeks post infection in aged mice (Fig. 6a), suggesting acute influenza viral pneumonia leads to chronic impairment of lung physiological function. Furthermore, increased lung resident T cells, which have a higher IFN-γ production capacity, could be observed in the aged mice post 6 weeks after infection (Fig. 6b, and Extended data Fig. 10a). To explore whether IFN-γ is important in the development of chronic lung sequelae in this model, we administered influenza-infected aged mice with IFN-γ neutralizing antibody following viral clearance at 14 dpi^49^ (Fig. 6c, and Extended data Fig. 10b). Anti-IFN-γ treated mice exhibited reduced lung and BAL cell counts including BAL T cells, indicating a potential IFN-γ-dependent feed forward loop for immune cell recruitment to the respiratory tract (Fig. 6d, and Extended data Fig. 10c). IFN-γ blockade also resulted in a pronounced reduction of both percentages and numbers of infiltrating monocytes in the respiratory tract (Fig. 6e, f, and Extended data Fig. 10d, e). Consistently, SiglecF^lo^ MoAM cells were significantly reduced (Fig. 6g).

Histopathological evaluation of the lungs highlighted a marked reduction in inflammatory lesions post IFN-γ blockade (Fig. 6h, i). Complementing this, Masson’s trichrome staining unveiled diminished collagen deposition following anti-IFN-γ intervention, indicating attenuated lung fibrosis (Fig. 6h). Previous reports have revealed R-PASC lung tissue with decreased alveolar type 1 (AT1) cells and AT2 cells^4^ ^50^. We found that AT1 staining densities (PDPN^+^) and AT2 cell counts (proSP-C^+^) are enhanced after IFN-γ neutralization (Fig. 6j, k), suggesting that IFN-γ blockade promoted lung functional recovery. Consequent assessments of lung compliance (Cst), respiratory damping (G), respiratory resistance (Rrs), and tissue elastance (Ers) affirmed a significant improvement of airway hyperresponsiveness, parenchyma injury and tissue fibrosis (Fig. 6l, and Extended data Fig. 10f). Collectively, our findings underscore that targeting IFN-γ offers a promising avenue to mitigate persistent lung sequelae and promote pulmonary functional recovery after acute viral pneumonia (Extended data Fig. 11).

## Discussion

Although a number of studies have associated dysregulated peripheral immune responses with PASC^10, 11, 12^, the immune landscape in a primarily affected organ, the lung, remains largely unknown in PASC. Here, integrating scRNAseq and clinical lung function evaluations, we noted a marked dysregulated MoAM cell responses associated with respiratory PASC. Pathway analysis underscored that respiratory resident T cell-derived IFN-γ drives MoAM precursor recruitment, polarization, and endorsing a pro-fibrotic property. Additionally, comparative analysis and functional studies in murine models of pulmonary sequelae established IFN-γ as a driver in chronic pathology and functional decline post-acute viral pneumonia (Extended data Fig. 11).

Notably, IFN-γ was associated with severe inflammation and alveolar injury during acute COVID-19^51^, and elevated serum IFN-γ levels^52^ were found in individuals with PASC after the resolution of primary infection^11^. However, those studies did not address whether IFN-γ acts as a “driver” or a “passenger” in the acute and chronic pathogenesis of SARS-CoV-2. Our comparative scRNAseq analysis coupled with functional neutralization at the post acute infection stage clearly established the role of IFN-γ in driving R-PASC, which likely further lead to systemic symptoms such as fatigue due to chronic hypoxia. Our analysis further unveiled two potential IFN-γ-mediated mechanisms in driving PASC, enhancement of inflammatory monocyte recruitment via the promotion of CCL2 production, and/or facilitating the development of pro-fibrotic MoAM polarization and differentiation. Additionally, IFN-γ has been recently shown to inhibit alveolar stem cell proliferation directly^53^, and thus it may inhibit the regeneration of alveolar and interstitial tissues through its direct signaling in alveolar epithelial cells following acute SARS-CoV-2 infection.

Our analysis revealed that lung-resident T cells were likely the major IFN-γ producing cells in PASC. Additionally, SARS-CoV-2 specific IFN-γ producing T cells were correlated with worse lung function, indicating aberrant virus-specific memory or “long-lived effector” T cells are a major culprit for pulmonary sequelae after acute COVID-19. Such a notion is consistent with previous findings by us and others^15, 18^. Of note, IFN-γ protein production by effector or memory T cell usually require concurrent TCR signaling. To this end, SARS-CoV-2 RNA can persist in various organs for weeks to months in humans^54, 55, 56^, and the duration of viral RNA shedding can last up to 59 days in the lower respiratory tract^57^. Therefore, it is possible that SARS-CoV-2 RNA or antigen persistence in the respiratory tract may drive the continued production of IFN-γ after the clearance of infectious virus^58^. Alternatively, chronic autoantigen release due to persistent tissue injury and/or innate inflammatory signals may also contribute to the production of IFN-γ in the respiratory tract^59^.

A few animal models with persistent tissue inflammation and fibrotic responses after acute SARS-CoV-2 infection have been developed. For instance, sustained pathology was first described in the hACE2-expressing humanized (MISTRG6-hACE2) mouse model^14^, although it is hard to assess the roles of dysregulated immune responses in this PASC model due to the lack of intact functional immune responses in those humanized mice. Recently, an aged BALB/c model of chronic disease after acute SARS-CoV-2 infection was developed and persistent sequelae including pulmonary lesions, immune infiltrate, and tissue fibrotic responses were observed in those infected mice^13^.

Interestingly, we found that the chronic tissue pathology observed in infected BALB/c mice is independent of IFN-γ as evidenced by the lack of the IFN-γ responsive signature in pulmonary cells and the inability of IFN-γ neutralization to ameliorate lung sequelae. This may be related to the bias toward Th2 responses in the BALB/c genetic background^60, 61, 62^. Therefore, it is crucial to perform a comparative analysis between human specimens and mouse PASC models to dissect the underlying molecular mechanisms driving PASC in patients for the development of relevant therapeutics in the future.

Through rigorous comparative analyses, we pinpointed a transient window in aged C57BL/6J mice that closely emulated the immune profile of R-PASC patients. Notably, the lung conditions in the R-PASC patients can persist for more than two years^2^. Given the short lifespan of rodents (one aged mouse day may be equal to 20 to 170 human days^63, 64^), the R-PASC window duration in aged C57BL6 mice in our model is perceivable. Interestingly, the recovery of the post SARS-CoV-2 sequelae at a later time point is linked to diminished IFN-γ signaling in the respiratory tract. Therefore, the brief window of persistent IFN-γ production and/or signaling may be the ultimate reason underlying the limited time window of R-PASC in mice. Thus, it is of interest to examine in the future whether forced extension of IFN-γ signaling by biologicals or genetic means could sustain pulmonary sequelae after SARS-CoV-2 infection. Furthermore, some patients do gradually recover from R-PASC over time^2^. Whether diminished IFN-γ signaling underlies the timely recovery of R-PACS requires future studies with kinetical BAL sampling, which is extremely difficult to perform given the invasive nature of BAL procedure in humans. Nevertheless, influenza viral pneumonia in aged mice appear to generate more persistent pulmonary sequelae^49^ ^50^, likely due to more persistent antigen deposit and/or wide spread of alveolar damage in the acute phase^65, 66^. IFN-γ neutralization did promote tissue regeneration and mitigate lung functional impairment in this prolonged lung sequelae model.

Thus, prolonged IFN-γ responses may be a common mechanism underlying the development of chronic pulmonary sequelae after acute respiratory viral infections. TRAMs typically maintain themselves through self-renewal independent of monocyte input during homeostasis. Under specific conditions such as infections or irradiation injuries, circulating monocytes are recruited to the alveolar space and can gradually adopt AM signature^45, 67, 68^. Notably, M2 polarized macrophages are generally considered as the macrophage subset mediating tissue fibrosis^69^. Studies have highlighted MoAMs in fibroblast foci within IPF lung tissues or after bleomycin injury, displaying a shift toward M2 phenotypes correlating with collagen deposition^70, 71, 72, 73, 74^. More recent reports also found that MoAMs may transiently upregulate both M1 and M2 genes during lung fibrosis development^20^. Nevertheless, our study connects IFN-γ-responsive M1-like MoAMs with tissue fibrosis development and lung functional impairment. This is likely due to the highly polarized Th1 responses in the lung microenvironments after viral infection. Future studies are needed to explore the potential underlying mechanisms by which those M1-like MoAMs cause tissue inflammation and fibrosis after acute SARS-CoV-2 infection.

The limited R-PASC cohort size (nine convalescents, two non-COVID-19 controls, and ten published healthy donor datasets, with a total of 85,971 BAL cells) restricts conclusive statements regarding specific patient comorbidities or treatment influences in this study. Nevertheless, through the evaluation of a published dataset featuring lung tissues from a distinct cohort of PASC pulmonary fibrosis patients ^4^, we were able to corroborate our observations across different cohorts. Notably, a larger cohort of R-PASC donors in a recent pre-print manuscript also supports our findings by identifying a population of MoAMs associated with persistent respiratory symptoms and radiographic abnormalities ^30^, although functional investigations with appropriate models were not performed in the study. Our research uniquely applied comparative analysis across human data and three different animal models to explore the molecular mechanisms underlying R-PASC development. Through these, we have revealed IFN-γ as a central node mediating exuberant T-monocyte-derived macrophage interactions, driving persistent pulmonary inflammation, tissue fibrosis and lung functional impairment in R-PASC. Our work emphasizes the necessity of performing a comparative analysis of human specimens and relevant animal-derived samples side-by-side to prob into the cellular and molecular origins of PASC for the development of future therapeutic tactics. Furthermore, our data strongly suggest that the JAK-inhibitor, Baricitinib, which has already been granted emergency use authorization for acute COVID-19 by United States Food and Drug Administration (FDA), may also serve as a promising candidate to treat ongoing respiratory PASC in the clinic.

## Acknowledgments

We thank Mayo genomic core, UVA Research Histology Core and Genome Analysis Technology Core for technical assistance. Cartoon in Extended data Fig. 10. was created with BioRender.com. We thank Dr. Maxim N. Artyomov and Dr. Sheng’en Hu during the data analysis. The study was in part supported by the US National Institutes of Health grants AI147394, AG069264, AI112844, AI176171 and AI154598 to J.S, and HL170961 to J.S. and R.V.

## Author contribution

C.L., W.Q., X.W. & J.S. conceived the overall project. C.L., W.Q., X.W., Y.W., H.N., M.A., I.S.C., K.S., R.K. and R.V. designed the experimental strategy and analyzed data, performed experiments, analyzed data, or contributed critical reagents to the study. C.L., W.Q., X.W. & J.S. wrote the original draft. All authors read, edited, and approved the final manuscript.

## Methods

### Ethics statement and biosafety

This study was approved by Mayo Clinic Institutional Review Board (protocol ID 20-004911). Informed consent was obtained from all enrolled individuals. All animal experiments were performed in animal housing facilities at the University of Virginia (UVA; Charlottesville, VA). The animal experiments were approved by UVA Institutional Animal Care and Use Committees. All work with SARS-CoV-2 infection was approved under Animal Biosafety Level 3/Biosafety Level 3 (A-BSL3/BSL3) conditions and was performed with approved standard operating procedures and safety conditions by the UVA Institutional Review Board.

### Study cohorts

Patients hospitalized due to PCR-confirmed SARS-CoV2 infection and COVID-19 pneumonia, requiring at least 2 liters of supplemental oxygen to manage respiratory failure, were considered convalescents of moderate to severe SARS-CoV-2 pneumonia. A control group of individuals matched for age and without lung disease were included in the study. When hospitalized patients needed more than 6 liters of supplemental oxygen, a high-flow nasal cannula (HFNC) oxygen delivery system was used, providing oxygen flows from 40 to 70 liters per minute, resulting in oxygen levels (FiO2) between 0.45 and 0.70 at the highest flow rate. The flow rate of the HFNC was adjusted at the bedside to maintain oxygen saturation between 88% and 92%.

The COVID-19 convalescent cohort consisted of patients between the ages of 60 and 85 with no evidence of preexisting interstitial or any prior chronic lung disease. For both the COVID-19 and control cohorts, previous lung disease was excluded through evaluation of the electronic medical records and clinical evaluation before performing a bronchoscopy. Patients with a history of <10 pack-years of smoking and mild chronic obstructive pulmonary disease (COPD) with FEV1 > 80% predicted and FEV1/FVC < 0.7 were still eligible for enrollment. At the time of bronchoscopy, control individuals had to have an absence of lung infiltrate, fever, or any signs of infection. Most controls underwent bronchoscopy to evaluate lung nodules or focal adenopathy of indeterminate cause. The exclusion criteria for both the COVID-19 and control cohorts included the inability to provide consent to participate in the study; patient under guardianship or curatorship; preexisting chronic lung disease including interstitial lung disease, pulmonary fibrosis, or any other chronic lung disease except for mild COPD as outlined in “inclusion criteria”; active cigarette smoking, vaping, or other inhalation use (former smoker providing quit >90 days before admission acceptable); and immunocompromised host status due to ongoing therapy with methotrexate, CellCept, azathioprine, prednisone dose >15 mg daily, rituximab, cyclophosphamide, or other immunosuppressive or other biologic agents. All patients with COVID-19 were enrolled for the study with bronchoscopy and BAL, acquisition of peripheral blood for PBMC, RNA research sample, peripheral blood for clinical preoperative clearance laboratories, and pulmonary function testing. The COVID-19 patient cohort was enrolled within a 60- to 90-day window after the onset of acute COVID-19 infection, the onset of which was defined as the day when the PCR SARS-CoV-2 swab was recorded as positive.

### Human pulmonary function tests

Pulmonary function testing was performed on COVID-19 and control cohort. All individuals underwent measurements of FVC and FEV1 as well as measurement of DLCO. In all individuals, spirometry was performed in the institutional pulmonary function laboratory at Mayo Clinic in accordance with American Thoracic Society (ATS) guidelines^75^.

### Bronchoscopy and BAL collection

Fiber-optic bronchoscopy and BAL were performed using moderate conscious sedation using standard clinical procedural guidelines in an outpatient bronchoscopy suite. Conscious sedation was administered in accordance with hospital policies, and a suitably trained registered nurse provided monitoring throughout the procedure. About 100 to 200 ml of saline were instilled in 20 ml aliquots until 60 ml of lavage fluid was obtained. The specimen was placed on ice, and immediately hand carried to the laboratory for analysis. The fluid collected was placed on ice and transferred immediately to the laboratory for processing.

### Single-cell purification from peripheral blood and BAL

The whole blood was mixed with phosphate-buffered saline (PBS) and then gently put over on Ficoll-Paque (Cytiva) in a 50-ml tube. Buffy coat generated by centrifuging at 400g for 40 min at RT was collected. For single-cell purification from BAL, BAL was filtered with a 70-μm cell strainer (Falcon) and then centrifuged at 300g for 10 min at 4°C. The supernatant was collected for multiplex assay and ELISA. The cells were collected for flow cytometry analysis and scRNAseq.

### Single-cell RNA sequencing library construction and results analysis

To facilitate the single-cell gene expression (GEX), PBMC and BAL cells from donors and MA10 infected mice at indicated time points were barcoded with 10X 5’ Library & Gel Bead Kit v1.1 or 3’ Library & Gel Bead Kit v3.1. A total of 10,000 cells were targeted for single-cell libraries preparation per the manufacturer’s instructions (10x Genomics).

scRNAseq were aligned and quantified using 10x Genomics Cell Ranger Software Suite (v4.0.0) against their corresponding human reference genome (GRCh38) downloaded from 10x Genomics website. Using default settings in Seurat 4.0.1 package, the filtered transcriptome data were then normalized (RNA expression by a factor of 10,000 with log-transformed). Aligned sequencing data were further analyzed with Seurat (v4.3.0). The threshold of percent.MT is 10 to exclude dead cells. The RNA expression data were then further scaled based on regressing the number of unique molecular identifiers (UMIs) detected and the percentage of gene counts per cell. Principal components analysis (PCA) was performed using the top variable genes. FindNeighbors and FindClusters functions were applied for cell clustering in Seurat for either dataset. Differential gene expression analysis was performed by the function of FindAllMarkers from Seurat with model-based analysis of single-cell transcriptomics (MAST) test or negbinom test, and gene set enrichment analysis (GSEA) analysis is based on the results of FindAllMarkers with the package of clusterProfiler^76^, and the online tool: Metascape^77^; AddModuleScore function was applied for analyzing cell population signature scores; Pseudotime analysis was performed using SeuratWrappers and Monocle 3 combination, based on the Seurat processed analysis at the single-cell level; transcription factor regulations were analyzed with SCENIC^31^. Additional BAL cells were extracted from the published dataset GSE151928^17^, and Aged-matched control PBMC scRNAseq data were merged with the public dataset^16^ (https://artyomovlab.wustl.edu/immune-aging/explore.html). Lung cell sc-RNA-seq data sets were adopted from: GSE224955, GSE146981 and GSE135893.

### Virus infection and antibody administration

Young female C57BL/6J mice were purchased from The Jackson Laboratory (JAX) at aged of 10 weeks and housed in specific pathogen-free animal facility for 2 weeks before infection. Aged female C57BL/6J mice were received at 20 to 24 months of age from the National Institute of Aging or the JAX and maintained in the same conditions for at least 1 month before infection. One-year-old female BALB/cJ mice were obtained from JAX at age of 10 weeks and aged to 1 year old. All mice were used under conditions fully reviewed and approved by the Institutional Animal Care and Use Committee guidelines at the University of Virginia.

Vero E6 cell line (ATCC CRL-1587) was maintained in Dulbecco modified Eagle medium (DMEM) supplemented with 10% fetal bovine serum (FBS), along with 1% of penicillin-streptomycin and L-glutamine at 37°C in 5% CO2. The SARS-CoV-2 mouse-adapted strain MA10 was kindly provided by Dr. Barbara J Mann (University of Virginia School of Medicine). MA10 was passed into Vero E6 cells and the titer was determined by plaque assay using Vero E6 cells^78^. Influenza A/PR8/34 virus stock was generated in our laboratory.

For SARS-CoV-2 MA10 infection in C57BL/6 mice, young or aged female C57BL/6J mice were intranasally infected with 5×10^4^ PFU MA10 virus. For SARS-CoV-2 MA10 infection in BALB/c mice, around 1-year-old female BALB/c mice were intranasally inoculated 10^3^ PFU SARS-CoV-2 MA10 virus as previously reported^13^. All control mice were intranasally inoculated with DMEM. For primary influenza virus infection, influenza A/PR8/34 strain was inoculated at the dose of ∼50 PFU as described before^49^. Infected mice were monitored daily for changes in weight and clinical signs of disease over a period of 2 weeks, following once a week throughout the duration of the experiments. The mortality rate of mice determined as “deceased” were either found dead in their cages or were euthanized upon reaching 70% of their initial body weight in accordance with the established humane endpoint specified in animal protocol. Mice were euthanized on days 0, 3, 10, 21 and 35 post infection. Bronchoalveolar lavage (BAL) samples were collected for flow cytometry and scRNAseq analysis. The lung lobes were preserved in 10% phosphate buffered formalin for 7 days prior being removed from BSL3 facility for further processing.

For in vivo blocking of IFN-γ, mice were injected intraperitoneally from 10 days post MA10 infection or 14 days post PR8 infection weekly, with 500μg (first dose) or 250μg (second and the following doses) of InVivoMAb anti-mouse IFNγ antibody (Clone: XMG1.2, Bio X Cell) or InVivoMAb rat IgG1 isotype control (Clone: HRPN, Bio X Cell).

### Mouse lung tissue dissociation and BAL cell collection

Mice were injected i.v. with 2 μg of anti-CD45 diluted in 200 μL of sterile PBS as previously described^79^. Mice were euthanized, and tissues were collected five minutes after injection of the i.v. Ab. Mouse BAL cells were obtained from BAL as described previously^80^, briefly, alveolar lavages were pooled from BAL washes (PBS with 2 mM EDTA). The lung tissues were harvested and dissociated in 37°C for 30 min with Gentle-MACS (Miltenyi). Single cell suspensions were further passed through 70μm cell strainers once before the next step operation as previously described^81^.

### Mouse lung tissue section and immunofluorescence staining

After euthanasia, mice were perfused with PBS (10 ml) via the right ventricle. Formalin (10%) was then gently instilled into the lung and left inflated for 1 min before excising and moving the lobe to 10% formalin for 48 hours followed by transfer to ethanol (70%). Samples were embedded in paraffin, and 5-μm sections were cut for hematoxylin, eosin (H&E), and Masson’s trichrome stain. Slides were then digitally scanned 20× resolution with the Aperio system (Leica), and the inflammatory area were quantified by QuPath software.

Immunofluorescence staining was performed on formalin-fixed paraffin-embedded (FFPE) lung tissue slides. FFPE slides were deparaffinized in Xyelene and later rehydrated prior to heat-induced antigen retrieval using 9pH Dako Target retrieval solution for 20 min. The slides were then blocked with 10% normal goat serum phosphate-buffered saline (PBS) for 30 min at room temperature (RT) and then were incubated with either Hamster anti-PDPN (Abcam Cat#: ab11936) or Rabbit anti-proSP-C (Millipore Sigma Cat#: AB3786) overnight at 4°C. After rinsing in 0.1% PBST (PBS with Tween 20) solution, the slides were incubated with Goat anti-Armenian Hamster IgG (H+L) (ThermoFisher Cat#: A78963) and Goat anti-rabbit IgG (H+L) (ThermoFisher Cat#: A-11036). Slides were aired after rinsing with 0.1% PBST before mounting with 4′,6-diamidino-2-phenylindole for nuclei counterstain. Tissue staining for the Ab mixture was reviewed, and images were captured using the Olympus BX63 microscope. For each lung section, images were taken in at least 10-12 random areas in the distal lung. All images were further processed and quantification by using ImageJ Fiji and/or QuPath software.

### Virus titer measurement

The viral titer in the BAL which collect from young and aged female C57BL/6J mice at 3 d.p.i, 10 d.p.i, 21 d.p.i and 35 d.p.i, were determined using plaque assay. Vero E6 cells were cultured in DMEM with the addition of 2% Fetal Clone II serum (Hyclone) and 1% Pen/Strep/glutamate. Serial dilutions were added to the cells. The plate was incubated at 37°C and 5% CO2 for 1 hour, shaking the plates every 15 minutes. After incubation, monolayers were overlayed with media containing 1.2% Avicel PH-101 and incubated at 37 °C and 5% CO2. After 72 hours, the overlay was removed, wells were fixed with 10% formaldehyde, and stained with 0.1% crystal violet to visualize plaques. Plaques were counted, and PFUs were calculated according to the following equation: Average # of plaques/dilution factor × volume diluted virus added to the well.

### Mouse lung function measurement

Lung function measurements using FOT and the resulting parameters have been previously described^49^. In brief, animals were anesthetized with an overdose of ketamine/xylazine (100 and 10mg/kg intraperitoneally) and tracheostomized with a blunt 18-gauge canula (typical resistance of 0.18 cmH2O s/mL), which was secured in place with a nylon suture. Animals were connected to the computer-controlled piston (SCIREQ flexiVent), and forced oscillation mechanics were performed under tidal breathing conditions described in^49^ with a positive-end expiratory pressure of 3 cm H2O. The measurements were repeated following thorough recruitment of closed airways (two maneuvers rapidly delivering TLC of air and sustaining the required pressure for several seconds, mimicking holding of a deep breath). Each animal’s basal conditions were normalized to their own maximal capacity. Measurement of these parameters before and after lung inflation allows for determination of large and small airway dysfunction under tidal (baseline) breathing conditions. Only measurements that satisfied the constant-phase model fits were used (>90% threshold determined by software). After this procedure, mice had a heart rate of ∼60 beats per minute, indicating that measurements were done on live individuals.

### Intracellular staining, antibodies, and flow cytometry

Cell suspensions were stained with indicated surface markers after Fc receptors blockade, staining was performed at 4 °C for 30 min. Cells were washed twice with FACS buffer (PBS, 2 mM EDTA, 2% FBS, 0.09% Sodium Azide), before fixation and permeabilization with either Perm Fix and Perm Wash (Biolegend, for cytokine staining) in the dark. Cells were washed twice with perm wash (BioLegend), stained with Abs for at least 30 min at RT and then washed twice with perm wash before flow cytometry acquisition. FACS Abs were primarily purchased from Biolegend, BD Biosciences or Thermo. The clone number of those Abs as follows: Zombie NIR, Anti-mouse-y6G (clone 1A8), Anti-mouse-Ly6C(clone HK1.4), Anti-mouse-SiglecF(clone E50-2440), Anti-mouse-CD8 (clone 53-6.7), Anti-mouse-CD4 (clone RM4-5), Anti-mouse-CD45 (clone 30-F11), Anti-mouse-CD11c (clone N418), Anti-mouse-CD11b (clone M1/70), Anti-mouse-MerTK (clone 2B10C42), Anti-mouse-CD64 (clone X54-5/7.1), Anti-mouse-MHCII (clone M5/114.15.2), Anti-mouse-CD44 (clone IM7), Anti-mouse-CD19 (clone SJ25C1), Anti-mouse-IFN-γ (clone XMG1.2), Anti-human-CD68 (clone Y1/82A), Anti-human-CD169 (clone 7-239), Anti-human-IFN-γ (clone 4S.B3), Anti-human-CD4 (clone OKT4), Anti-human-CD8a (clone RPA-T8). The dilution of surface staining Abs was 1:200 and dilution of intracellular staining Abs was 1:100. After staining, cells were acquired through an 14-color Attune NXT system (Life Technologies). Data were then analyzed by FlowJo software (Treestar).

### Bone marrow and circulating monocytes derived macrophage like cell differentiation and stimulation

C57BL/6J mice derived Bone marrow cells or human blood derived monocytes (purified with CD14 MicroBeads from PBMC) were harvested and treated as describe^40^, briefly, recombinant mouse or human GM-CSF (20 ng/ml), mouse or human TGF-β (2 ng/ml), and 0.1 μM PPAR-γ agonist rosiglitazone were added into the culture system, 8 days after differentiation, differentiated cells were treated with recombinant mouse or human IFN-γ for 24 hours. Culture medium and RNA were collected for the following experiments.

### Statistical analysis

To compare between two sample groups, Mann-Whitney test was applied for unpaired comparisons. For analysis between several groups, one- or two-way analysis of variance (ANOVA) was performed; Wilcoxon rank-sum test and MAST were performed during scRNAseq data analysis. Correlations were assessed by the Pearson correlation coefficient running under R in RStudio (1.4). All statistical tests were performed using GraphPad Prism version 10 (GraphPad Software) or R (version 4.0.3).

## Data and materials availability

scRNAseq data are available from the Gene Expression Omnibus under accession number. All other data needed to evaluate the conclusions of the paper are present in the paper or the Supplementary Materials.

---

**Extended Data Figure 1.**
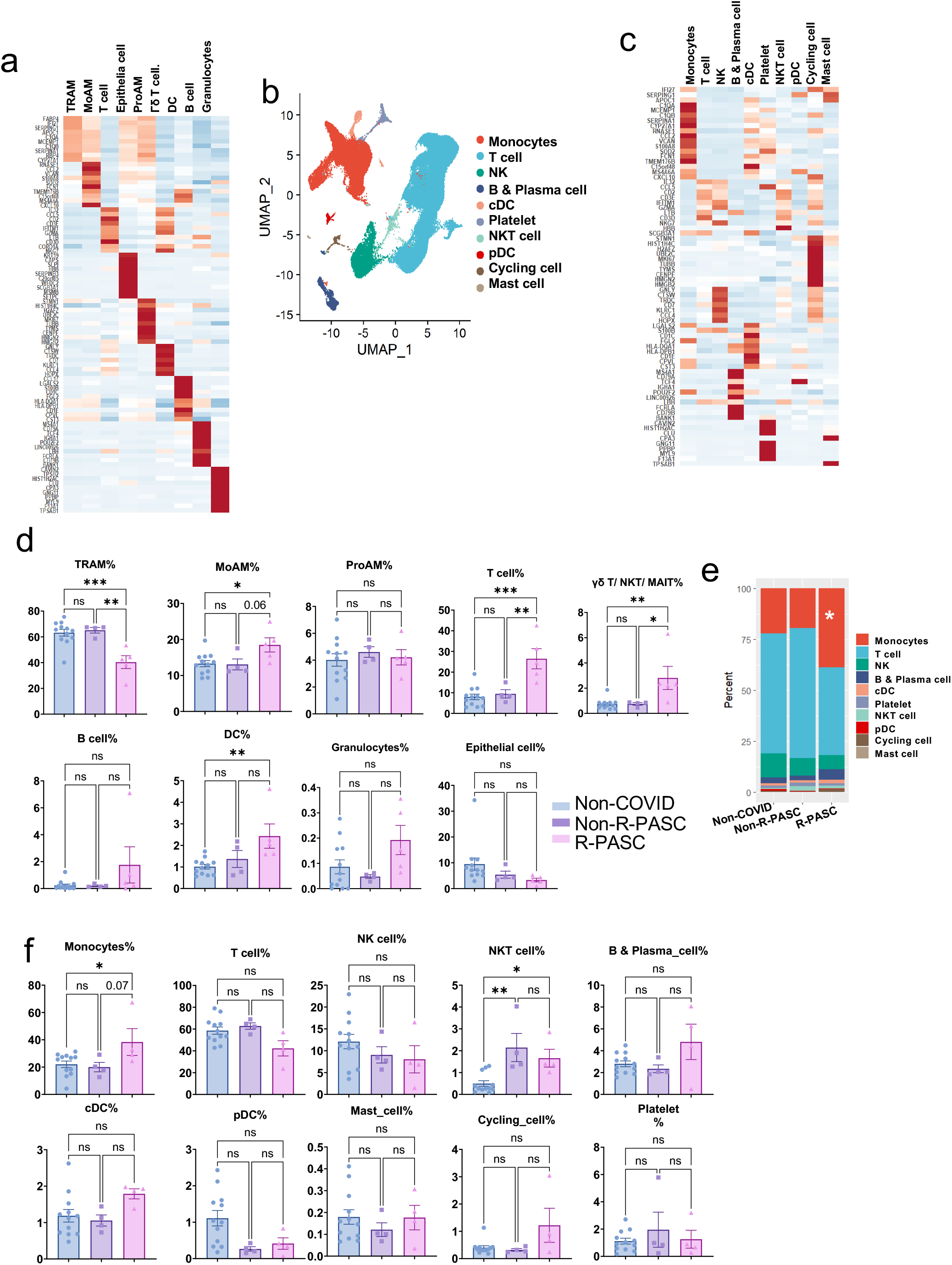
R-PASC patients exhibit an altered immune cell composition. **a**, The top 10 representative genes of each cell cluster from BAL. **b**, The UMAP plot of integrated analysis of PBMCs from donors. **c**, The top10 representative genes of each cell cluster from PBMCs. **d**, Proportions of cell clusters in BAL among non-COVID, non-R-PASC and R-PASC groups. **e**, Stacked bar plots showing the proportion of indicated cell types in PBMCs among non-COVID, non-R-PASC and PASC groups. **f**, Proportions of cell clusters in PBMC among non-COVID, non-R-PASC and R-PASC groups. Data are represented mean ± SEM unless otherwise indicated. Significance were tested by one-way ANOVA with Tukey’s adjustment for multiple comparisons, ∗p < 0.05; ∗∗p < 0.01; ∗∗∗p < 0.001; ∗∗∗∗p < 0.0001.

**Extended Data Figure 2.**
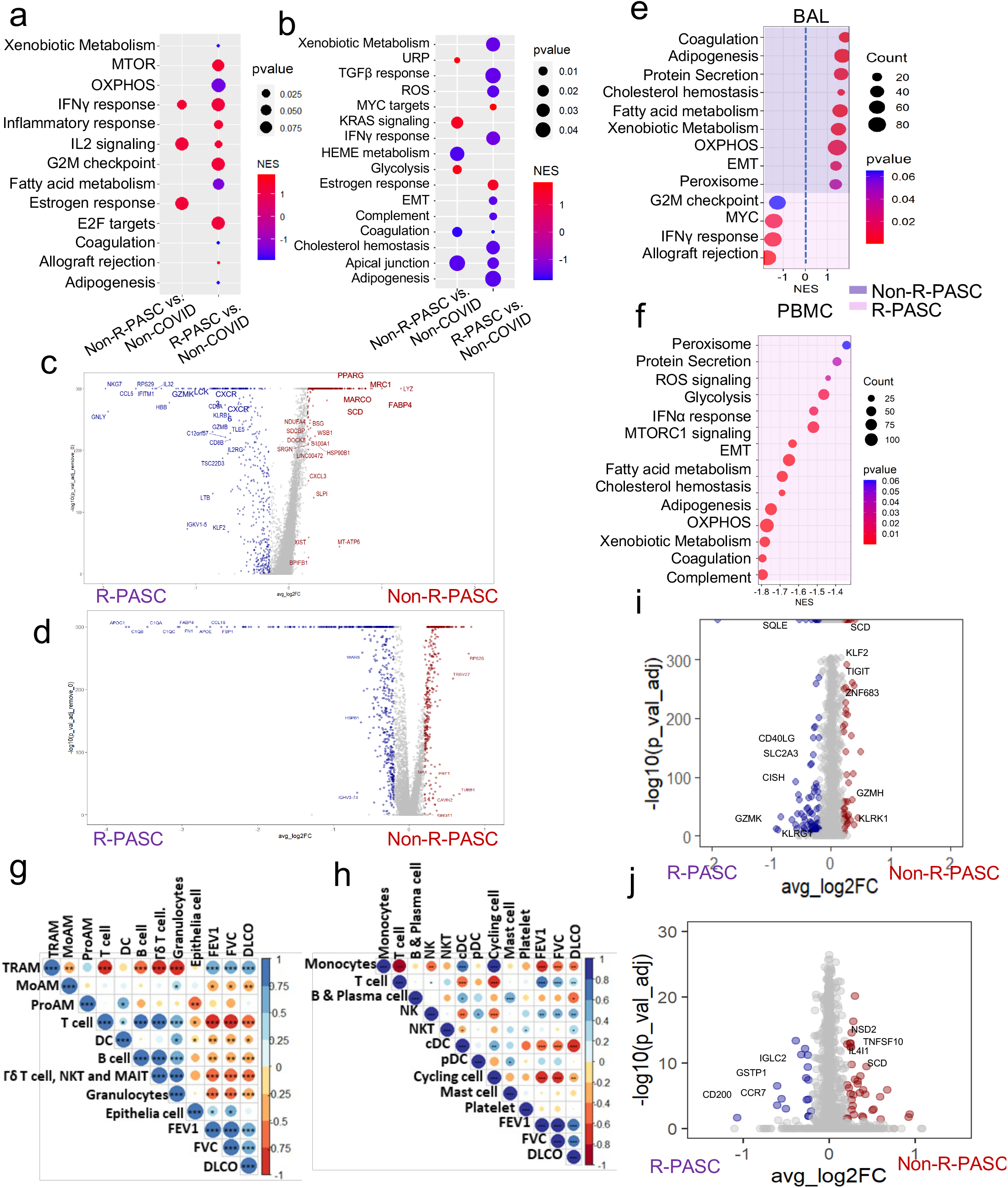

R-PASC patients exhibit an altered immune landscape.

**a**, **b**, Dot plots showing the BAL cells (**a**) and PBMCs (**b**) enriched pathways in non-R-PASC or R-PASC groups compared to non-COVID controls.

**c, d,** Volcano plots showing the difference expressed genes in BAL cells (**c**) and PBMCs (**d**) between non-R-PASC and R-PASC donors.

**e, f,** GSEA results of BAL cells (**e**) and PBMCs (**f**) from non-R-PASC or R-PASC donors.

**g**, **h**, Correlation of lung functional parameters with properties of BAL cells (**g**) or PBMCs (**h**).

**i, j,** Different expressed genes of BAL T cells (**i**) and BAL B cells (**j**) between non-R-PASC and R-PASC donors.

∗p < 0.05; ∗∗p < 0.01; ∗∗∗p < 0.001; ∗∗∗∗p < 0.0001.

**Extended Data Figure 3.**
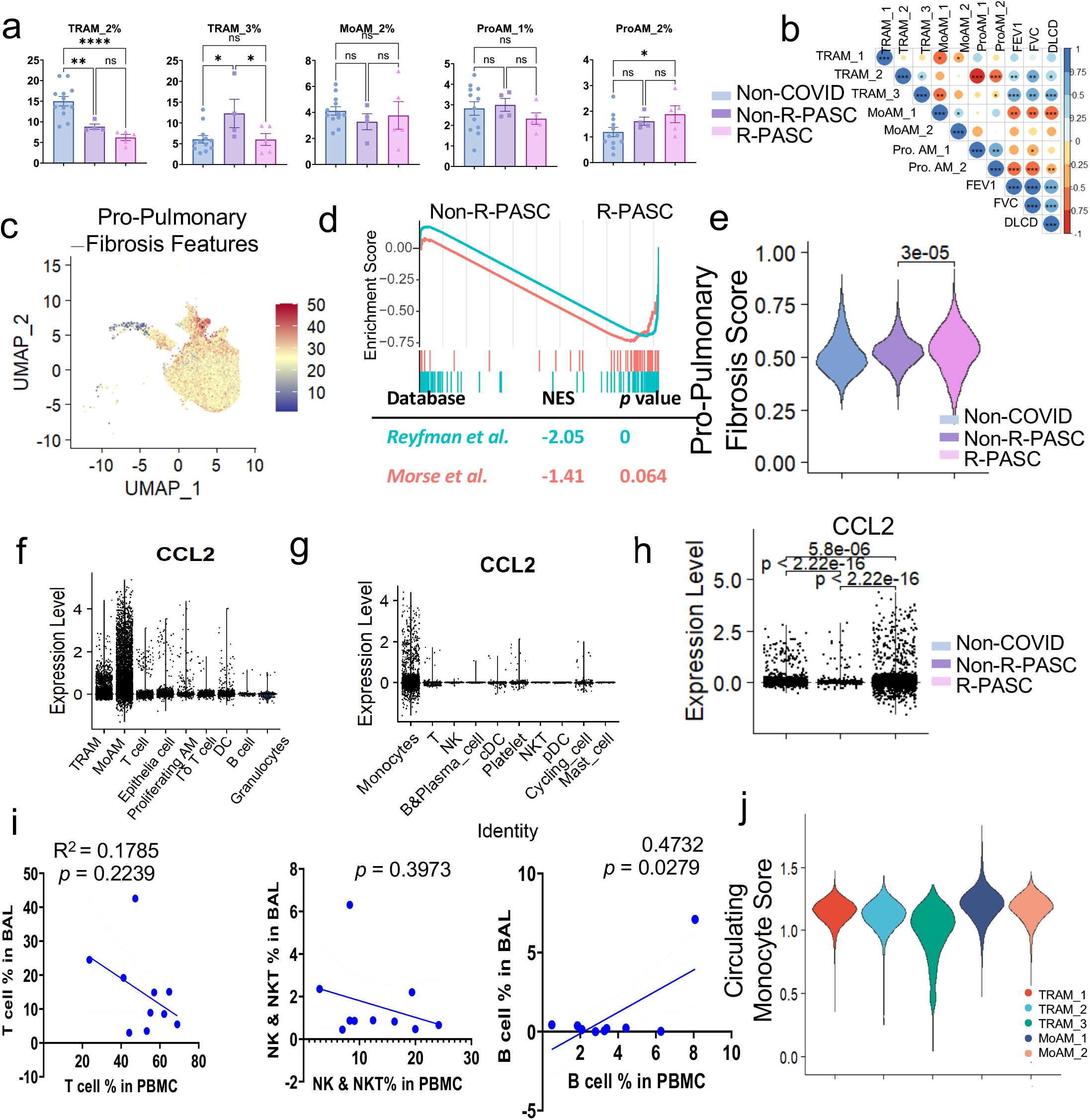
Respiratory macrophage phenotypes in R-PASC individuals. **a,** Percentage of BAL macrophage clusters among non-COVID, non-R-PASC and R-PASC groups. **b**, Correlation of lung function parameters and BAL macrophage clusters proportion. **c**, Feature plot of pro-pulmonary fibrotic macrophage gene signature. **d**, GSEA plots of pulmonary fibrosis related macrophage gene sets between non-R-PASC and R-PASC BAL Macrophages. **e**, Relative scores of a pro-pulmonary fibrotic macrophage gene signature in MoAM_1 cells from indicated groups. **f – h**, Violin plots showing the *CCL2* expression level in BAL cells (**f**), PBMCs (**g**), and BAL MoAM_1 cells (**h**) in indicated groups. **i**, Correlation of BAL T cells, NK and NKT cells, and B cells percentages with their circulating counterparts. **j**, PBMC monocyte feature assessment in BAL macrophage clusters. Data are represented mean ± SEM unless otherwise indicated. Significance were tested by one-way ANOVA with Tukey’s adjustment or Wilcoxon test, ∗p < 0.05; ∗∗p < 0.01; ∗∗∗p < 0.001; ∗∗∗∗p < 0.0001.

**Extended Data Figure 4.**
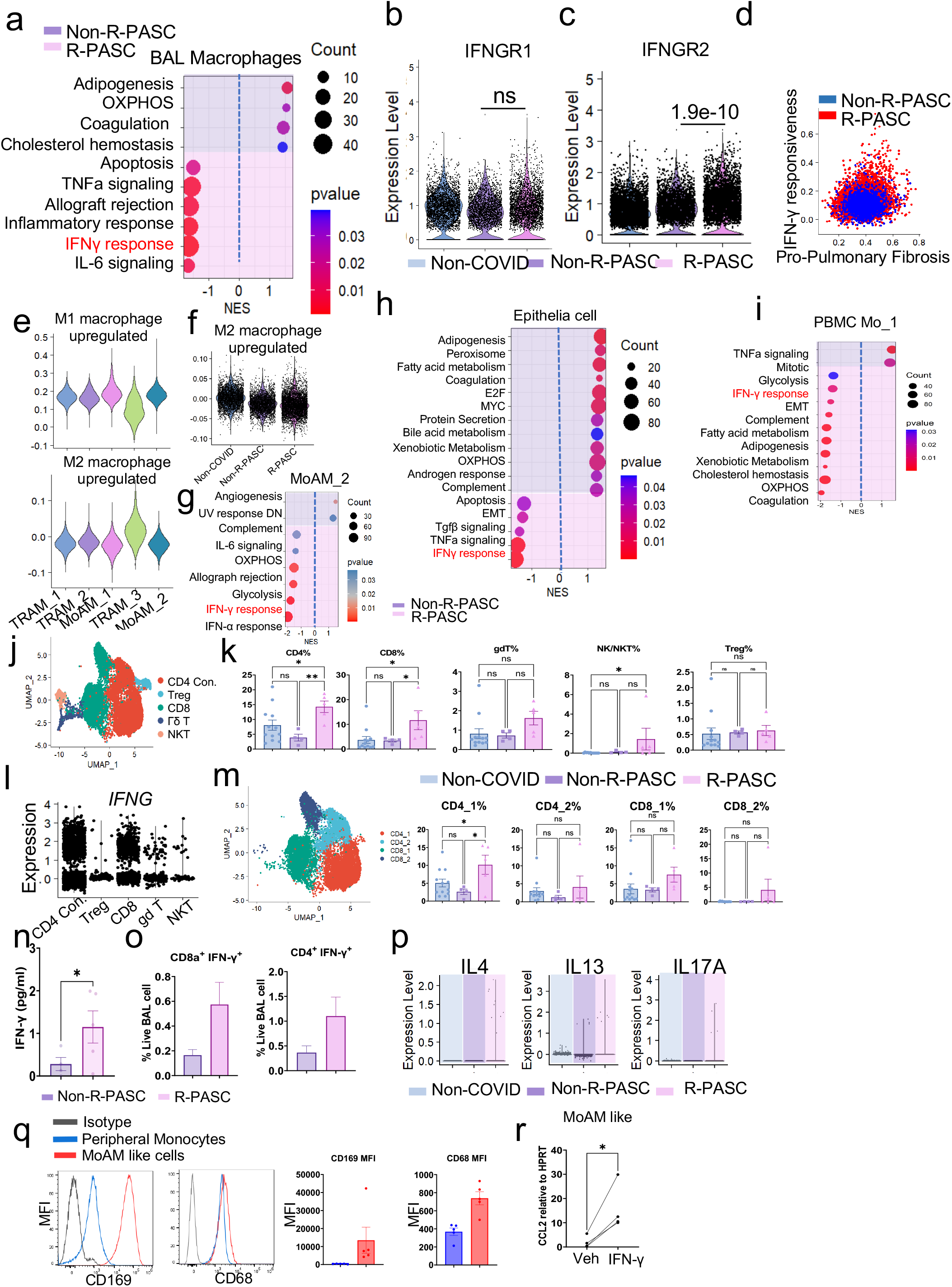
IFN-**γ** as a critical node mediating exuberant T-macrophage interactions in R-PASC. **a**, Gene sets enriched in the BAL macrophages from Non-R-PASC or R-PASC donors. **b, c**, Violin plot showing the *IFNGR1* (**c**) and *IFNGR2* (**d**) expression in the MoAM_1 cell from indicated groups. **d**, Scatter plot showing the IFN-γ responsiveness score and pro-fibrotic score in non-R-PASC and R-PASC MoAM_1cells. **e**, M1 (upper) or M2 (lower) macrophage features among the BAL macrophage clusters. **f**, M2 macrophage feature score of non-COVID, no-R-PASC and R-PASC MoAM_1cells. **g-i**, Gene sets enriched in the BAL MoAM_2 cluster (**g**), epithelial cell cluster (**h**), and PBMC Monocyte_1 cluster (**i**) from Non-R-PASC or R-PASC donors. **j**, The UMAP plot of BAL T cells. **k**, Proportions of T cell clusters in BAL among non-COVID, non-R-PASC and R-PASC. **l**, *IFNG* expression in BAL T cell subpopulations. **m**, UMAP plot of BAL CD4^+^ conventional T and CD8^+^ T cells subpopulations(left). Proportions of BAL CD4^+^ conventional T and CD8^+^ T cell subpopulations among non-COVID, non-R-PASC and R-PASC (right). **n,** IFN-γ concentration in the BALF from indicated groups tested by multiplex assays. **o,** IFN-γ^+^ CD8^+^ or CD4^+^ T cell percentage of whole BAL cells in response to SARS-CoV-2 peptides stimulation. **p**, Violin plot showing the *IL4*, *IL13* and *IL17A* mRNA expression in the T cells from indicated groups. **q**, MFI of CD169 and CD68 in the peripheral blood derived monocytes or peripheral blood derived monocyte differentiated alveolar macrophage like cells. **r,** *CCL2* mRMA expression in the MoAM like cells post Vehicle or recombinant IFN-γ treatment. Data are represented mean ± SEM unless otherwise indicated. Significance were tested by one-way ANOVA with Tukey’s adjustment for multiple comparisons or Wilcoxon test, ∗p < 0.05; ∗∗p < 0.01; ∗∗∗p < 0.001; ∗∗∗∗p < 0.0001.

**Extended Data Figure 5.**
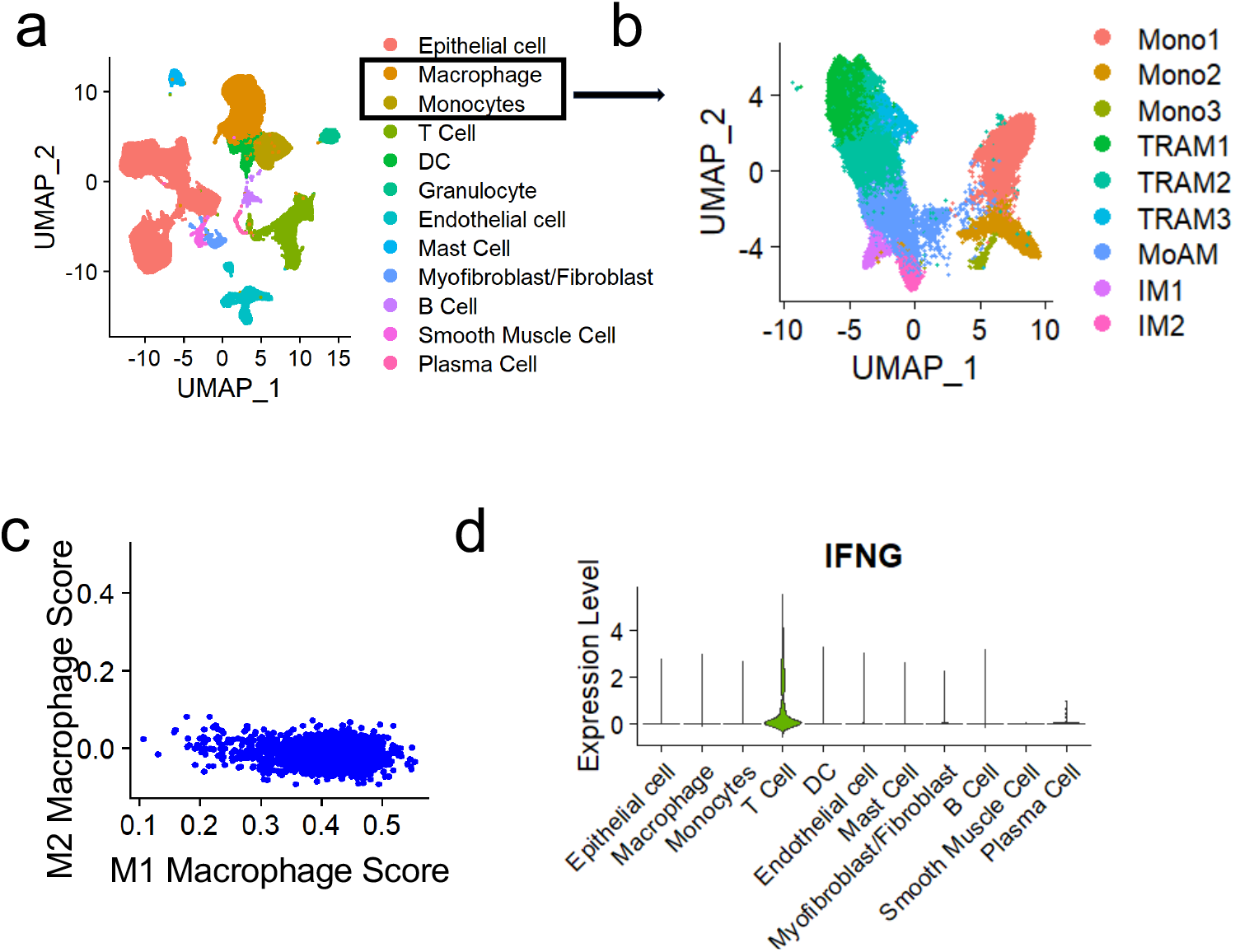
sc-RNA-seq analysis of PASC pulmonary fibrosis (PASC-PF) lung tissues. **a,** The integrated UMAP of cells from Non-COVID control or PASC-PF patients’ lung tissues ^4, 41, 42^. **b,** The integrated UMAP of macrophages from Non-COVID control or PASC-PF patients’ lung tissues. **c,** Scatter plots showing and M1 and M2 macrophage features in MoAM cells from lung tissues. **d,** *IFNG* expression in indicated cell types from lung tissues.

**Extended Data Figure 6.**
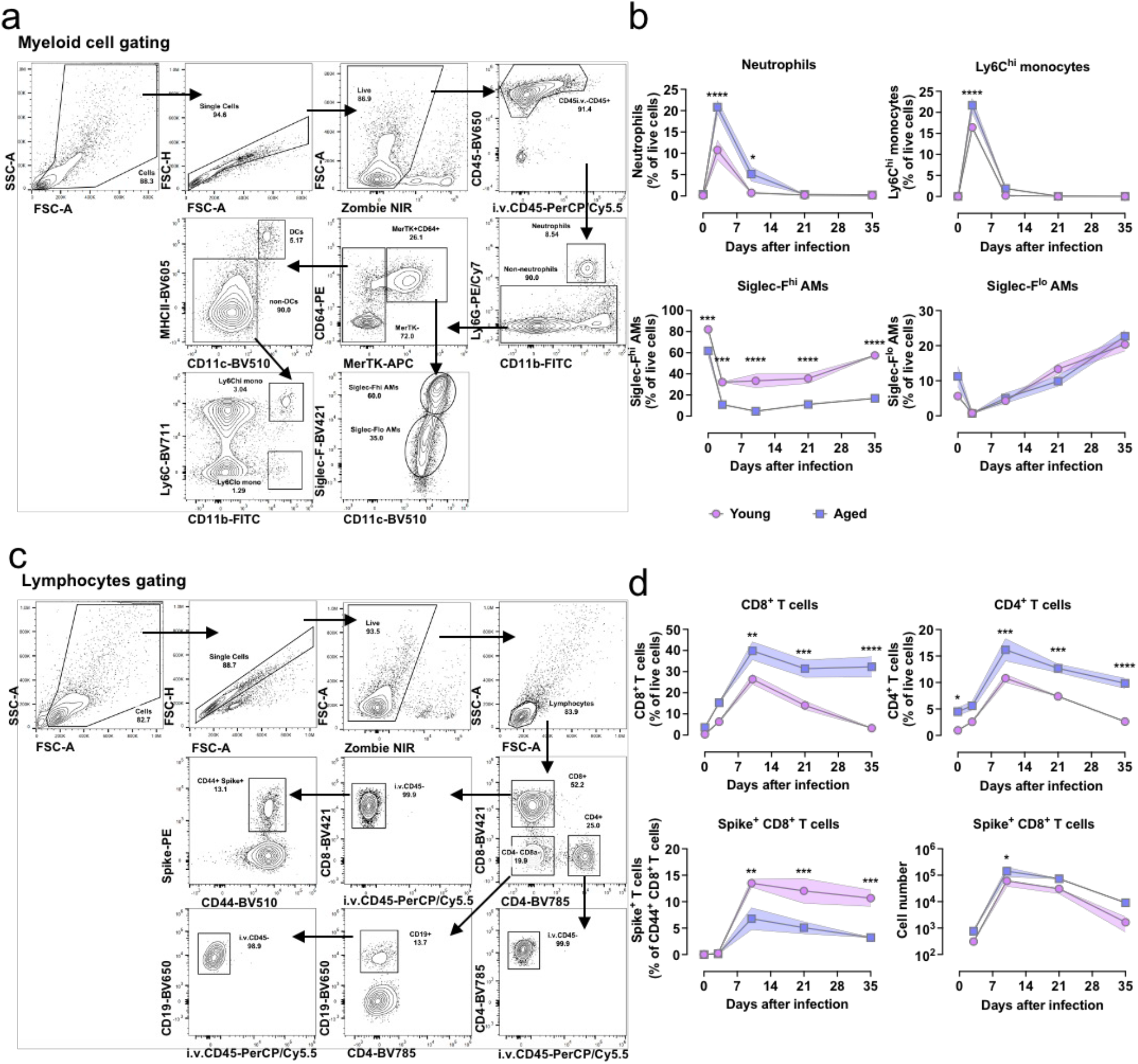
Flow cytometry gating strategy and percentages of major immune cells in BAL of mice infected with MA10. **a**, Representative flow cytometry plots of gating strategy for myeloid cells in BAL of young and aged mice infected with MA10. **b**, The percentage of neutrophils, Ly6C^hi^ monocytes, Siglec-F^hi^ TRAMs, and Siglec-F^lo^ MoAMs. **c**, Representative flow cytometry plots of gating strategy for T cells and Spike-specific CD8^+^ T cells in BAL of MA10-infected young and aged mice. **d**, The percentage of CD8^+^ T cells, CD4^+^ T cells and Spike-specific CD8^+^ T cells, as well as the number of Spike-specific CD8+ T cells. Data are represented mean ± SEM, and were analyzed by two-way ANOVA with Tukey’s adjustment for multiple comparisons, * p < 0.05, ** p < 0.01, *** p < 0.001, and **** p < 0.0001.

**Extended Data Figure 7.**
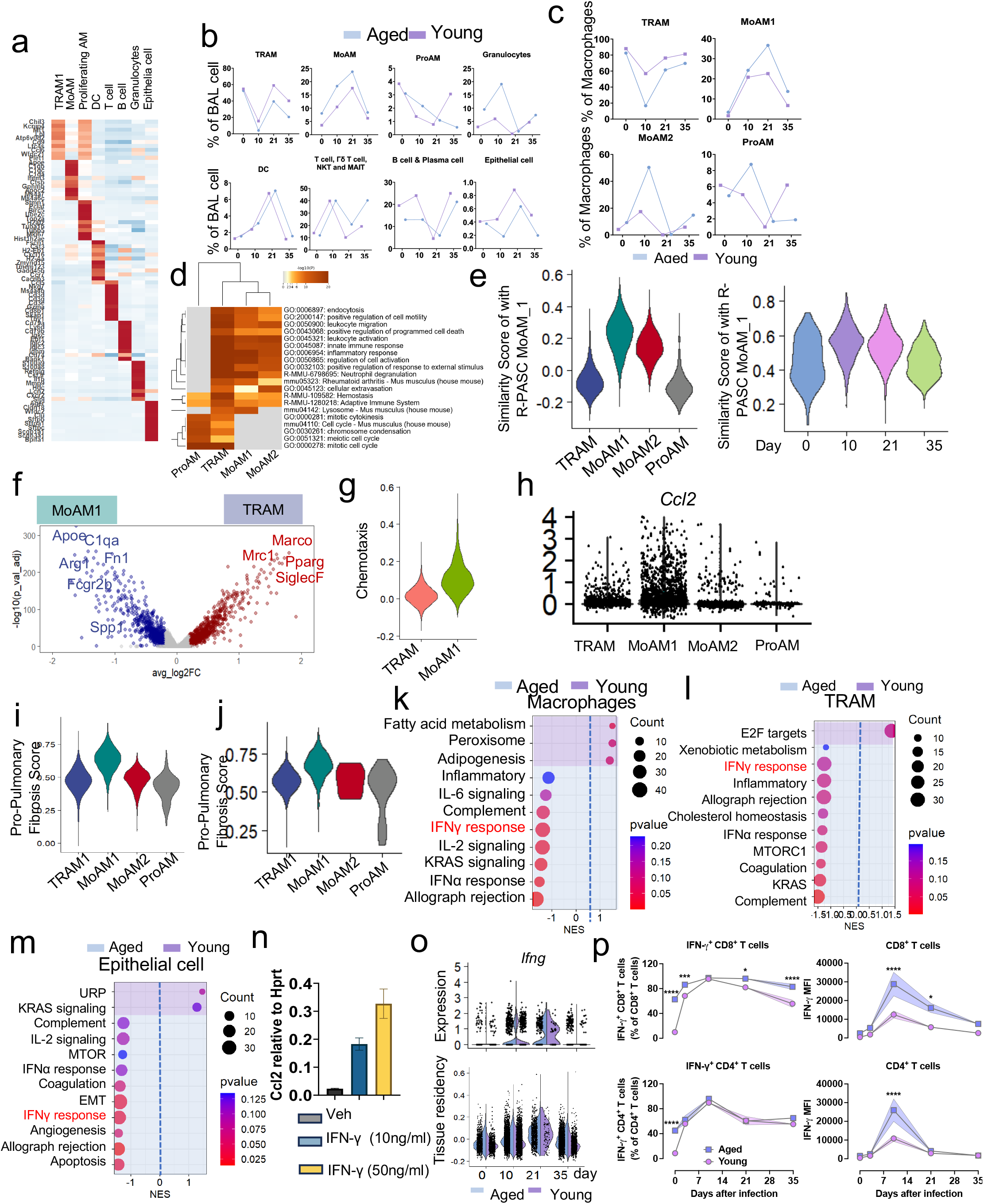
sc-RNA-seq analysis of BAL cells of a mouse R-PASC model. **a**, The top10 representative genes of each cell cluster from integrated C57BL/6J mice BAL. **b,** The proportion of each cell type in MA10 infected young and aged C57BL/6J mice at indicated time points. **c,** The proportion of each BAL macrophage subclusters in MA10 infected young and aged C57BL/6J mice at indicated time points. **d,** Heatmap of BAL macrophage subclusters enriched pathways. **e,** The similarity score of human MoAM_1 features in mouse BAL macrophage subclusters (**left**) or in MoAM1 cells at indicated time points (**right**) **f,** Volcano plot showing differentially expressed genes between MoAM1 and TRAM at 21 days post MA10 infection. **g**, Chemotaxis ability of MoAM1 and TRAM at 21 days post MA10 infection. **h,** *Ccl2* expression level in the BAL macrophage subclusters. **I, j,** Relative score of pro-pulmonary fibrotic macrophage gene signature in BAL macrophage clusters from all time points (**i**), or at day 21 post MA10 infection (**j**). **k - m,** Enriched gene sets of BAL macrophages (**k**), epithelial cell (**l**) and TRAM clusters (**m**) from young or aged C57BL/6J mice at 21 days post M10 infection. **n,** Relative expression of *Ccl2* in bone marrow derived MoAM like cells exposure to IFN-γ. **o,** *Ifng* expression (**upper**) and tissue residency score (**lower**) in BAL T cells from young or aged C57BL/6J mice at indicated time points post MA10 Infection. **p,** The percentage of IFN-γ-producing CD8^+^ T and CD4^+^ T cells, as well as MFI of IFN-γ in CD8^+^ T and CD4^+^ T cells, from young and aged mice infected with MA10 at indicated time points. Data represent the mean ± SEM. Data were analyzed by two-way ANOVA, * p < 0.05, ** p < 0.01, *** p < 0.001, and **** p < 0.0001.

**Extended Data Figure 8.**
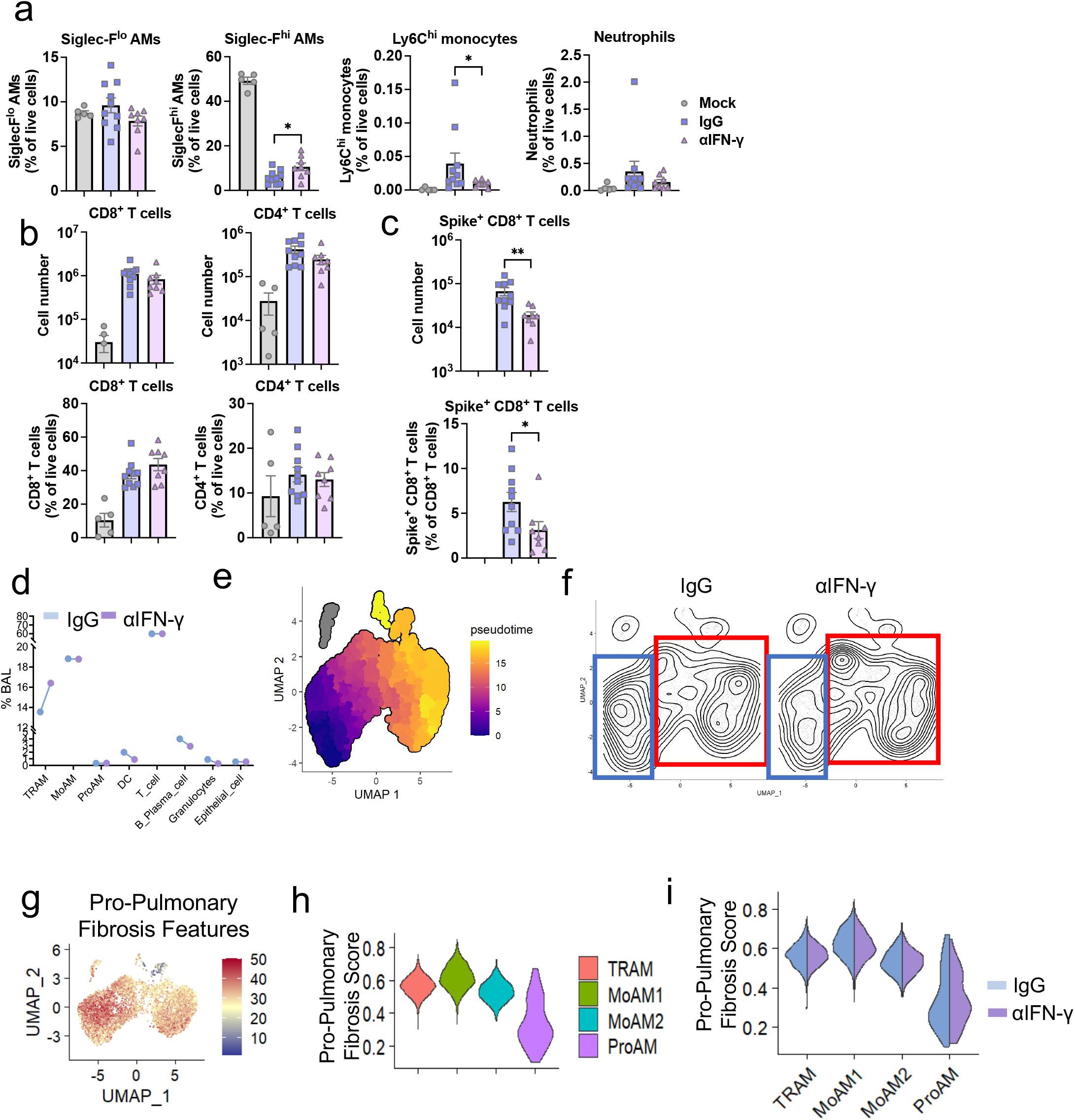
Targeting IFN-γ ameliorates post SARS-CoV-2 lung sequelae. **a,** The proportion of Siglec-F^lo^ AMs, Siglec-F^hi^ AMs, Ly6C^hi^ monocytes, and neutrophils in BAL of MA10-infected aged mice treated with IgG or αIFN-γ. **b, c,** The cell counts and percentage of CD8^+^ T and CD4^+^ T cells (**b**) and Spike^+^ CD8^+^ T cells (**c**)from aged mice treated with IgG or αIFN-γ. **d,** The proportion of indicated BAL cell types from IgG or αIFN-γ treated aged mice. **e**, Pseudotime analysis of BAL macrophage differentiation. **f**, Contour plots showing the density of BAL macrophage clusters in the UMAP. **g,** Feature plot of pro-pulmonary fibrotic features in BAL macrophages. **h, i,** Relative score of pro-pulmonary fibrotic macrophage gene signature in all BAL macrophage clusters (**h**), or in indicated groups (**i**). Data represent the mean ± SEM. Data were analyzed by one-way ANOVA or unpaired *t* test, * p < 0.05, ** p < 0.01, *** p < 0.001, and **** p < 0.0001.

**Extended Data Figure 9.**
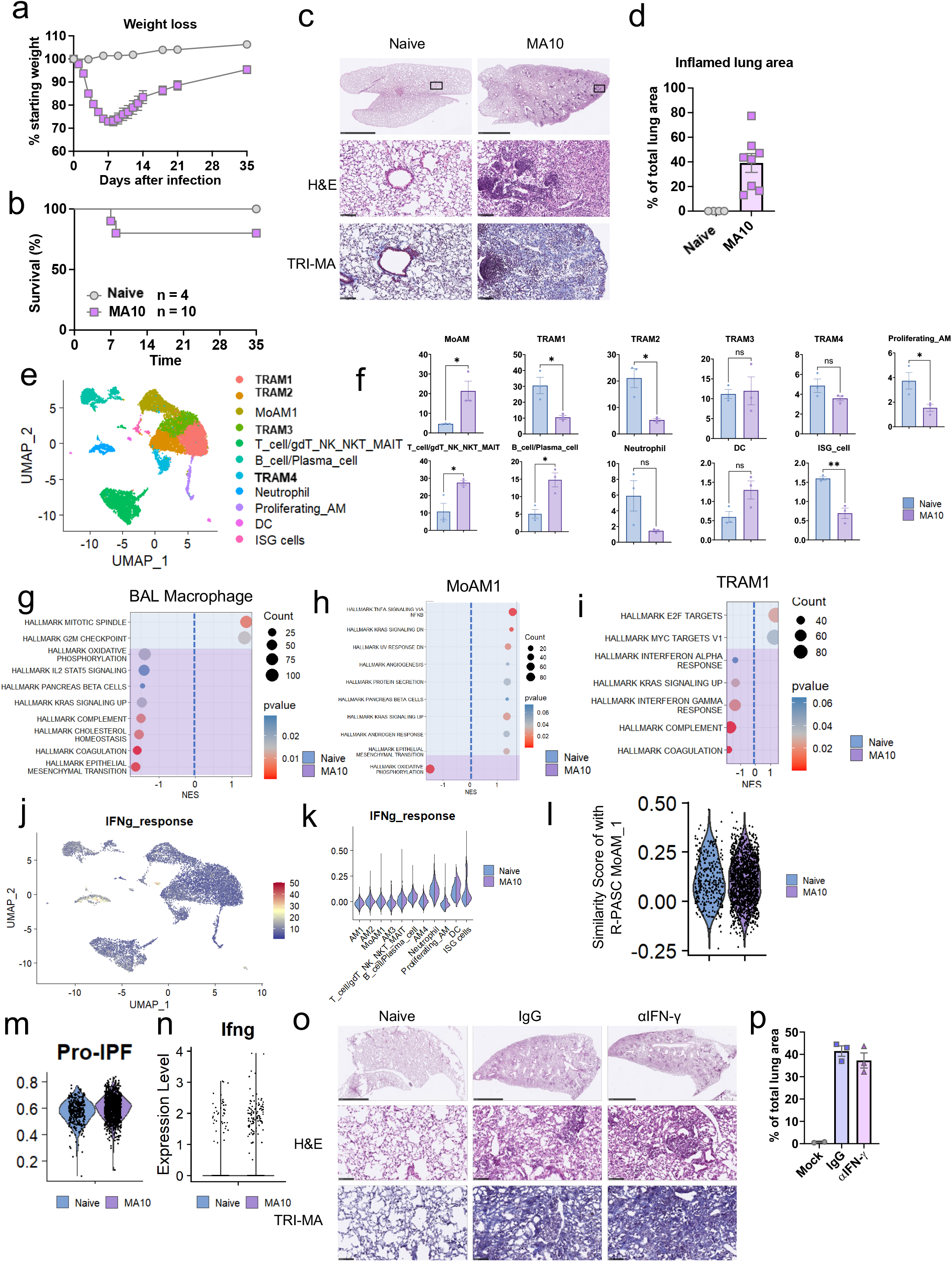
IFN-γ response is dispensable for post SARS-CoV-2 sequelae in BALB/c mice. **a** and **b,** Weight loss (**a**) and survival (**b**) of aged BALB/c mice were monitored post infection. **c,** Representative images of histopathology are shown. H&E indicates hematoxylin and eosin staining. Masson’s trichrome (TRI-MA) staining highlights fibrotic collagen deposition. Scale bar indicates 2.5 mm for whole lung lobe or 100 mm for zoomed area. **d**, QuPath quantification of inflamed area in the lungs for indicated time points as in **c**, represented as the percentage of total lung area. **e,** The integrated UMAP of BAL cell from naïve or 35 days post MA10 infected aged BALB/c mice, pooled from 3 mice each group. **f,** The proportion of each cell type in naïve or 35 days post MA10 infected aged BALB/c mice. **g-i,** Enriched gene sets of BAL macrophages (**g**), MoAM1 clusters (**h**) and TRAM1 clusters (**i**) from aged BALB/c mice at 0 or 35 days post MA10 infection. **j,** INF-γ responsiveness features in BAL cell from aged BALB/c mice at day 0 and day 35. **k,** Relative score of INF-γ responsiveness in indicated cell types from naïve or 35 days post MA10 infected aged BALB/c mice. **l,** The similarity score of human MoAM_1 feature in MoAM1 cells from indicated groups. **m,** Relative score of pro-pulmonary fibrotic macrophage gene signature in MoAM1 cells from indicated groups. **n,** Violin plot showing the *Ifng* expression in the T cells from indicated groups. **o,** Representative images of histopathology in the αIFN-γ or IgG treated aged BALB/c mice at 30 days post MA10 infection. H&E indicates hematoxylin and eosin staining. Masson’s trichrome (TRI-MA) staining highlights fibrotic collagen deposition. Scale bar indicates 2.5 mm for whole lung lobe or 100 mm for zoomed area. **p** QuPath quantification of inflamed area in the lungs for indicated treatment groups as in **o**, represented as the percentage of total lung area. Data represent the mean ± SEM. Data were analyzed by unpaired *t* test, * p < 0.05, ** p < 0.01, *** p < 0.001, and **** p < 0.0001.

**Extended Data Figure 10.**
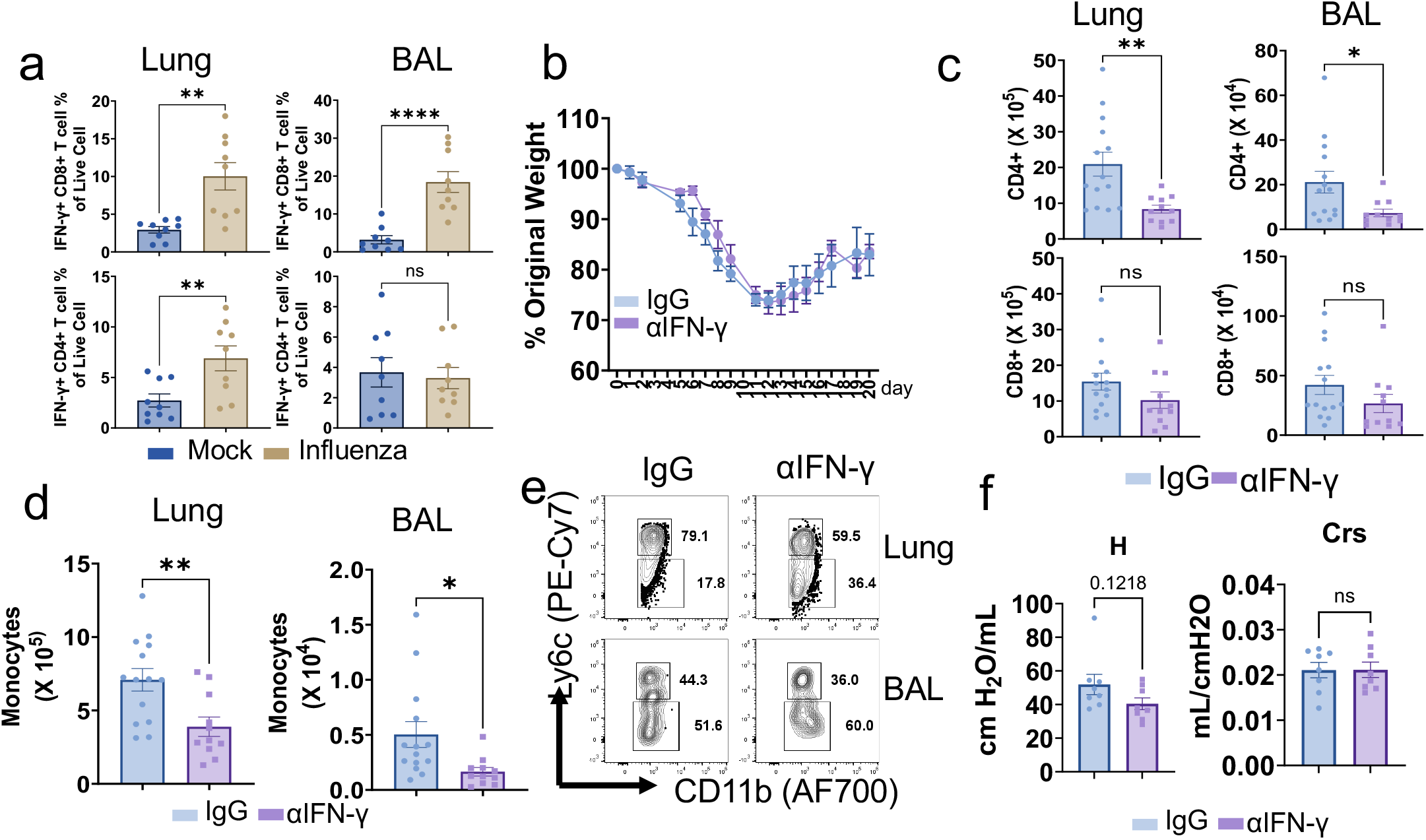
IFN-γ blockade promotes lung functional recovery in a persistent lung sequelae model after influenza pneumonia. **a,** IFN-γ producing lung resident (left) and BAL (right) CD8^+^ and CD4^+^ T cell proportion in naïve or influenza infected aged C57BL/6J mice at 6 weeks post infection, n= 9, pooled from two independent experiments. **b,** Weight loss of the infected aged mice post treatment, n= 4-6. **c,** Tissue resident CD4^+^ T cell and CD8^+^ T cell counts in lung (left) and BAL (right) post treatment, n= 11-14, pooled from three independent experiments. **d,** Monocytes counts in lung (left) and BAL (right) post treatment. **e,** Representative dot plots of Ly6c^hi^ or Ly6C^lo^ monocytes in lung (left) and BAL (right) post treatment. **f,** Evaluation of respiratory compliance (Crs), tissue resistance (H) post treatment with flexiVent, n= 8, pooled from two independent experiments. Data represent the mean ± SEM. Data were analyzed by unpaired *t* test, * p < 0.05, ** p < 0.01, *** p < 0.001, and **** p < 0.0001.

**Extended Data Figure 11.**
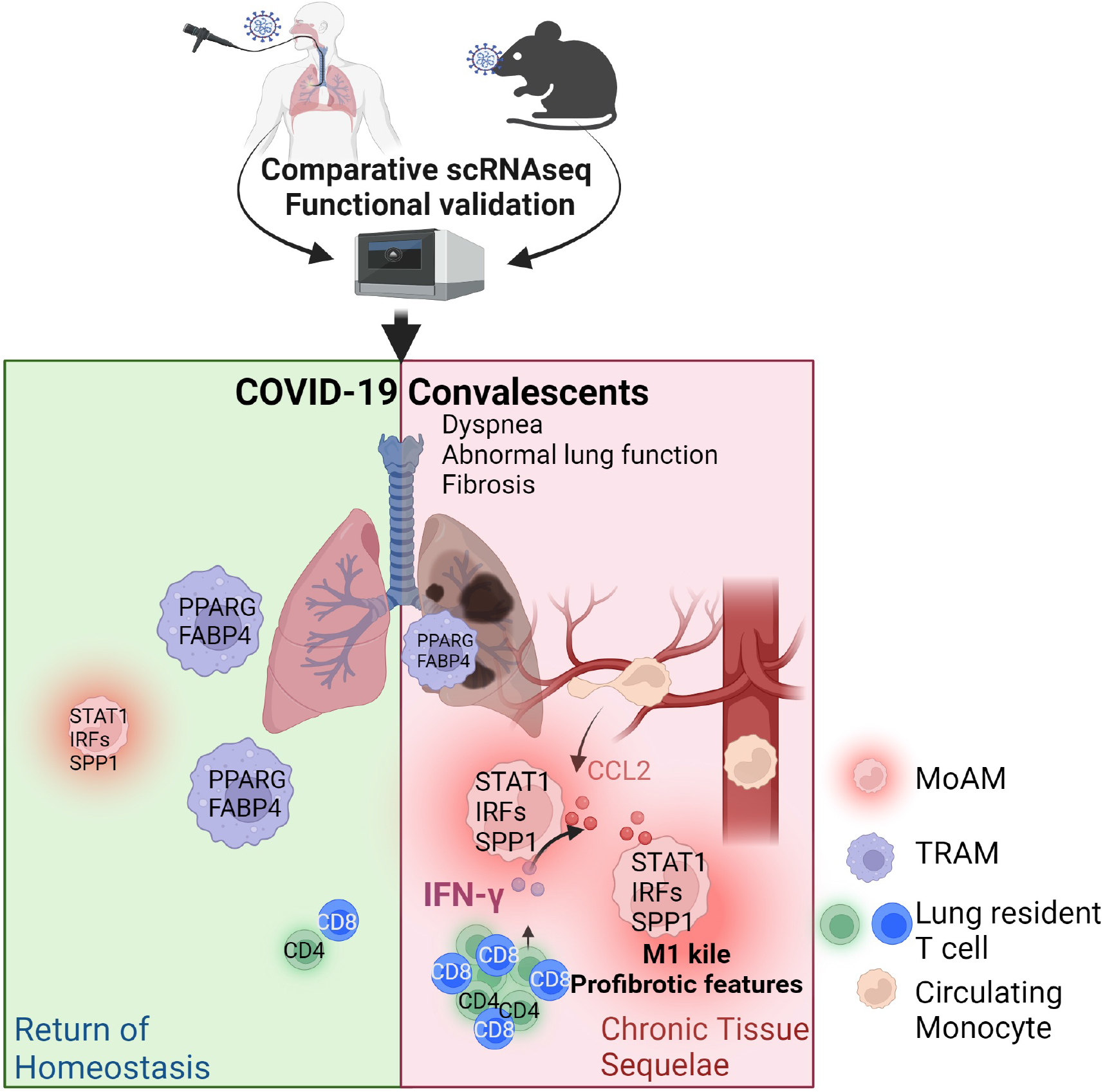
Graphic model.

Patient description and Pulmonary Function Test.

Patient description and Pulmonary Function Test
| Subject ID | Gender | AGE | Hospitalization Following COVID-19 diagnosis | Maximum O2 need (L/min) | ICU Stay (days) / Mechanical Ventilation (days) | Pulmonary Embolism | Secondary Bacterial Pneumonia | Other complications (type) | Date of bronchoscopy following COVID-19 diagnosis | FEV1 | FVC | DLCO | Grouping | sc-RNA-seq |
| --- | --- | --- | --- | --- | --- | --- | --- | --- | --- | --- | --- | --- | --- | --- |
| CON-1 | M | 65 |  |  |  |  |  |  |  | 112 / 3.54 / 2.33 | 116 / 4.80 / 3.10 | 60 / 15.1 / 18.6 | Non-COVID | No |
| CON-3 | F | 77 |  |  |  |  |  |  |  | 124 / 2.35 / 1.35 | 124 / 3.07 / 1.76 | 89 / 15.9 / 13.4 | Non-COVID | Yes |
| CON-4 | M | 73 |  |  |  |  |  |  |  | 114 / 3.16 / 1.98 | 116 / 4.24 / 2.69 | 109 / 24.9 / 16.7 | Non-COVID | Yes |
| CON-5 | M | 73 |  |  |  |  |  |  |  | 119 / 3.30 / 1.97 | 126 / 4.59 / 2.68 | 72 / 16.4 / 16.4 | Non-COVID | No |
| CVD-01 | M | 64 | D 5-14<br>D19-22 | 12 L | 7 / - | Yes | No | No | D 74 | 72 / 2.54 / 2.62 | 70 / 3.23 / 3.49 | 56 / 15.5 / 20.7 | R-PASC | Yes |
| CVD-02 | M | 83 | D10-18 | 3 L | - / - | No | No | No | D 74 | 85 / 2.14 / 1.73 | 76 / 2.60 / 2.44 | 75 / 16.3 / 15.7 | R-PASC | Yes |
| CVD-03 | M | 63 | D 5-26 | HFNC 60L | 11 / CPAP | Yes | No | No | D 74 | 68 / 2.39 / 2.61 | 64 / 2.95 / 3.46 | 45 / 12.5 / 20.5 | R-PASC | Yes |
| CVD-04 | M | 65 | D 3-14 | HFNC 60L | 8 / + 5 | No | Yes | No | D 88 | 81 / 2.42 / 2.20 | 75 / 2.88 / 2.90 | 92 / 21.8 / 17.6 | R-PASC | No |
| CVD-05 | M | 74 | D 6-16 | HFNC 60L | 3 / - | Yes | No | No | D 74 | 78 / 2.08 / 1.90 | 69 / 2.45 / 2.59 | 46 / 10.2 / 16.2 | R-PASC | No |
| CVD-06 | F | 66 | D 5-10 | 5 L | - / - | No | No | No | D 87 | 127 / 3.18 / 1.85 | 129 / 4.17 / 2.40 | 87 / 18.8 / 16.2 | Non-R-PASC | Yes |
| CVD-07 | M | 62 | D 5-10 | 2 L | - / - | No | No | No | D 84 | 112 / 4.06 / 2.70 | 99 / 4.66 / 3.58 | 99 / 28.0 / 21.1 | Non-R-PASC | Yes |
| CVD-08 | F | 76 | D 9-23 | HFNC 40L | - / - | No | No | No | D 80 | 83 / 1.64 / 1.40 | 77 / 1.99 / 1.83 | 44 / 8.2 / 13.9 | R-PASC | Yes |
| CVD-09 | F | 63 | D 6-14 | HFNC 60 L | 5 / - | No | No | No | D 78 | 53 / 1.36 / 1.77 | 52 / 1.69 / 2.26 | 60 / 11.9 / 14.9 | R-PASC | Yes |
| CVD-10 | M | 68 | D 1 - 13 | HFNC 50 L | 3 / - | No | No | No | D 67 | 119 / 4.09 / 2.50 | 114 / 5.17 / 3.39 | 81 / 22.2 / 20.3 | Non-R-PASC | Yes |
| CVD-11 | F | 64 | D2 - 21 | 50% FIO2 | 11/+9 | No | No | No | D 64 | 93 / 4 / 2 | 87 / 1 / 2.4 | 73 / 18.8 / 14.2 | Non-R-PASC | Yes |

